# Microbial magnetite oxidation via MtoAB porin-multiheme cytochrome complex in *Sideroxydans lithotrophicus* ES-1

**DOI:** 10.1101/2024.09.20.614158

**Authors:** Jessica L. Keffer, Nanqing Zhou, Danielle D. Rushworth, Yanbao Yu, Clara S. Chan

## Abstract

Most of Earth’s iron is mineral-bound, but it is unclear how and to what extent iron-oxidizing microbes can use solid minerals as electron donors. A prime candidate for studying mineral-oxidizing growth and pathways is *Sideroxydans lithotrophicus* ES-1, a robust, facultative iron oxidizer with multiple possible iron oxidation mechanisms. These include Cyc2 and Mto pathways plus other multiheme cytochromes and cupredoxins, and so we posit that the mechanisms may correspond to different Fe(II) sources. Here, *S. lithotrophicus* ES-1 was grown on dissolved Fe(II)-citrate and magnetite. *S. lithotrophicus* ES-1 oxidized all dissolved Fe^2+^ released from magnetite, and continued to build biomass when only solid Fe(II) remained, suggesting it can utilize magnetite as a solid electron donor. Quantitative proteomic analyses of *S. lithotrophicus* ES-1 grown on these substrates revealed global proteome remodeling in response to electron donor and growth state and uncovered potential proteins and metabolic pathways involved in the oxidation of solid magnetite. While the Cyc2 iron oxidases were highly expressed on both dissolved and solid substrates, MtoA was only detected during growth on solid magnetite, suggesting this protein helps catalyze oxidation of solid minerals in *S. lithotrophicus* ES-1. A set of cupredoxin domain-containing proteins were also specifically expressed during solid iron oxidation. This work demonstrated the iron oxidizer *S. lithotrophicus* ES-1 utilized additional extracellular electron transfer pathways when growing on solid mineral electron donors compared to dissolved Fe(II).

**Importance:** Mineral-bound iron could be a vast source of energy to iron-oxidizing bacteria, but there is limited physiological evidence of this metabolism, and it has been unknown whether the mechanisms of solid and dissolved Fe(II) oxidation are distinct. In iron-reducing bacteria, multiheme cytochromes can facilitate iron mineral reduction, and here, we link a multiheme cytochrome-based pathway to mineral oxidation, expanding the known functionality of multiheme cytochromes. Given the growing recognition of microbial oxidation of minerals and cathodes, increasing our understanding of these mechanisms will allow us to recognize and trace the activities of mineral-oxidizing microbes. This work shows how solid iron minerals can promote microbial growth, which if widespread, could be a major agent of geologic weathering and mineral-fueled nutrient cycling in sediments, aquifers, and rock-hosted environments.

## Introduction

To microbes, minerals provide surfaces to live on, a source of nutrients, and in some cases, a substrate for respiration, e.g. for Fe(III)- and S(0)-reducing organisms. We are increasingly finding that microbes can also oxidize minerals, particularly iron minerals such as magnetite (1–3), green rust (4), pyrite (5), biotite (6), and smectites (6–8), using these as a source of electrons, and therefore energy. To use minerals as electron donors, cells must be able to transfer electrons from outside the cell to the interior. This capability, known as extracellular electron uptake (EEU) has been demonstrated not only in cultures with minerals but also by experiments on cathodes in bioelectrochemical systems, which provide a continuous supply of electrons directly to colonizing cells (9–13). EEU is a capability of iron-oxidizing bacteria (FeOB), which need to keep iron outside of cells to prevent various detrimental reactions from occurring in the periplasm or cytoplasm (14, 15). Most work on FeOB has focused on oxidation of dissolved Fe^2+^, but if this EEU capability can be adapted to oxidize solid minerals, it would give an energetic advantage, given that most of Earth’s iron is mineral-bound.

However, we do not know how common mineral oxidation is amongst microorganisms. To recognize and track mineral oxidation, we need to unravel the mechanisms, i.e. the genes and proteins involved. This requires an organism that can grow both on dissolved and solid substrates. Among the few reliable chemolithotrophic FeOB isolates, the Gallionellaceae *Sideroxydans lithotrophicus* ES-1 stands out as having a versatile metabolism, able to grow by oxidizing dissolved Fe^2+^, Fe(II)-smectite clays, as well as thiosulfate (7, 16–18). *Sideroxydans* species have been identified in many environments, including a variety of sediments (19, 20), brackish, freshwater, or groundwater systems (16, 21–30), and rice paddies or other wetlands (31–33), suggesting this genus is highly adaptable, likely linked to its metabolic versatility.

*S. lithotrophicus* ES-1 has a closed, sequenced genome that encodes multiple possible enzymatic pathways for iron oxidation (17, 34, 35). The genome encodes three isoforms of the iron oxidase Cyc2, a fused monoheme cytochrome-porin which has been biochemically demonstrated to oxidize dissolved Fe^2+^ (36, 37). It also encodes MtoAB which is homologous to the iron-oxidizing PioAB complex of the photoferrotroph *Rhodopseudomonas palustris* TIE-1 (38) and the iron-reducing complex MtrAB in *Shewanella* species (39, 40). Porin-cytochrome complexes like MtrAB and PioAB form conduits across the outer membrane, so are key in iron-reducer interactions with minerals (MtrAB; (41, 42)), and in oxidation of a cathode (PioAB; (43)). Given the predicted structural similarity to these other systems, and that heterologously expressed MtoA has been shown to oxidize Fe(II) (44), MtoAB could also play a role in oxidation of extracellular minerals. The genome of *S. lithotrophicus* ES-1 also encodes other porin-cytochrome complexes with large multiheme cytochrome subunits and a plethora of heme motif (CXXCH)-containing proteins including probable periplasmic electron carriers (34, 45). Thus, *S. lithotrophicus* ES-1 appears well-endowed with multiple potential iron oxidation and other EEU mechanisms, though it is not certain which ones enable oxidation of minerals.

Recent work on *S. lithotrophicus* ES-1 demonstrated for the first time the ability of this organism to utilize a solid Fe(II) source for growth, and gave us some initial clues to the possible mineral oxidation mechanism (7). The porin MtoB was detected in cells grown on Fe(II)-smectite clays but not dissolved Fe(II)-citrate. The multiheme cytochrome MtoA was not observed, possibly because multiheme cytochromes can be difficult to detect by mass spectrometry due to the large number of covalently modified cysteines per peptide length. The proteomics was supplemented with RT-qPCR, which confirmed that *mtoA* was upregulated on smectite compared to Fe(II)-citrate. This led to the hypothesis that in *S. lithotrophicus* ES-1, the MtoAB complex plays a specific role in oxidation of solid iron minerals, but not aqueous Fe(II)-citrate (7). However, given that only a limited proportion of proteins (<25% of total proteome) were detected in this study, improvements to enhance proteome coverage for low-input samples are necessary to accurately distinguish proteins expressed on solid substrates.

Incomplete proteomes can result from low biomass input, as can often be the case for FeOB, since cultures are challenging. In the smectite study of *S. lithotrophicus* ES-1, large volumes of cultures were required to obtain enough cells for molecular analyses such as proteomics (7). Recently, this need for large culture volumes was eliminated with the development of a novel on-filter in-cell (OFIC) processing pipeline for proteomic analyses of low biomass samples (46–48). This single-vessel method avoids cell lysis, which tends to cause significant sample loss particularly for low-input samples and performs all the treatments in the same filter device, thus drastically simplifies sample preparation and improves proteomic sensitivity. In a pilot study, ∼76% of the entire *S. lithotrophicus* ES-1 proteome was identified from just ten milliliters of culture (∼1×10^9^ cells) (46).

Minerals with high Fe(II) content commonly interfere with molecular extractions, making it difficult to obtain complete ‘omics’ datasets. In the smectite study, clays interfered with downstream analyses (7), so we investigated the possibility of using magnetite, which can be easily removed from cultures with a magnet. As a mixed-valence iron mineral (Fe^II^Fe^III^_2_O_4_) common in sediments (49), magnetite could potentially serve as an electron donor to support the growth of Fe(II)-oxidizing bacteria. We hypothesized *S. lithotrophicus* ES-1 could grow by oxidizing Fe(II) in magnetite, in part because *S. lithotrophicus* ES-1 grows on other iron minerals, and also based on previous observations of other FeOB that were able to oxidize magnetite. The photoferrotroph *Rhodopseudomonas palustris* TIE-1 oxidized chemically synthesized magnetite (1, 50) while nitrate-reducing Fe(II)-oxidizers including *Acidovorax* sp. 2AN and the enrichment culture KS have been observed to oxidize biogenic magnetite (2, 3). If *S. lithotrophicus* ES-1 is able to oxidize magnetite, this would give us an optimal system for investigating proteins involved in solid Fe(II) oxidation.

Here, we tested *S. lithotrophicus* ES-1 growth on three batches of abiogenic magnetite (two synthesized in house and one purchased from a commercial vendor) and compared protein expression to cells grown on dissolved Fe^2+^. The substrates differed in particle size, crystallinity, and solubility, which allowed us to evaluate growth and Fe(II) oxidation mechanisms in the presence of different proportions of solid and dissolved Fe^2+^. This work gives further evidence that FeOB can grow by oxidizing mineral-bound Fe(II) along with insight into the mechanisms that enable electron uptake from solids.

## Results

### Magnetite characterization

We characterized the magnetites to determine particle size, crystallinity and solubility. The X-ray diffraction (XRD) patterns of fresh synthetic magnetite, aged synthetic magnetite, and commercial magnetite all possessed peaks characteristic of magnetite (Fig. S1). The sharp, narrow peaks in the commercial magnetite XRD pattern indicate the particles are more crystalline, and the particle size is calculated to be ∼27 nm. The synthetic magnetites have broader peaks in their XRD patterns, indicating lower crystallinity/smaller domain size, with the fresh synthetic magnetite having the smallest size (<10 nm).

Nanocrystalline minerals tend to be more soluble (51, 52) and this was confirmed by suspending 1 g/L (12.9 mM Fe) of magnetite particles in anoxic 20 mM MES buffer (pH 6.0) and measuring dissolved Fe^2+^ over the course of 24 hours (Fig. 1). The fresh synthetic magnetite was the most soluble, releasing a maximum of 570 μM Fe^2+^ (∼13% of total Fe(II)), which fits with the lower crystallinity of this phase. The aged synthetic magnetite was less soluble, releasing at most 179 μM Fe^2+^ (∼4% of total Fe(II)) and the commercial magnetite was the least soluble at <10 μM Fe^2+^ (the limit of detection in the assay; <0.2% of total Fe(II)). To estimate the dissolved Fe^2+^ released from the synthetic magnetites over a longer time in the absence of cells, dissolved Fe^2+^ concentrations were measured at 24-hour intervals in incubations using either anoxic buffer or buffer equilibrated with 2% oxygen to simulate the conditions for culturing. The buffer was then replaced with fresh solution to remove all dissolved Fe^2+^, and dissolved Fe^2+^ was re-measured after an additional 24 hours, and the process repeated once more. Each day, the dissolved Fe^2+^ release decreases, implying there is less soluble Fe(II) available over time. By the third incubation, the dissolved Fe^2+^ released from the fresh and aged synthetic magnetites was <100 μM (Fig. S2). Having magnetites of different solubilities allows us to evaluate growth and mineral oxidation mechanisms in the presence of different amounts of dissolved Fe^2+^, covering a range of possible environmental scenarios.

**Figure 1.**
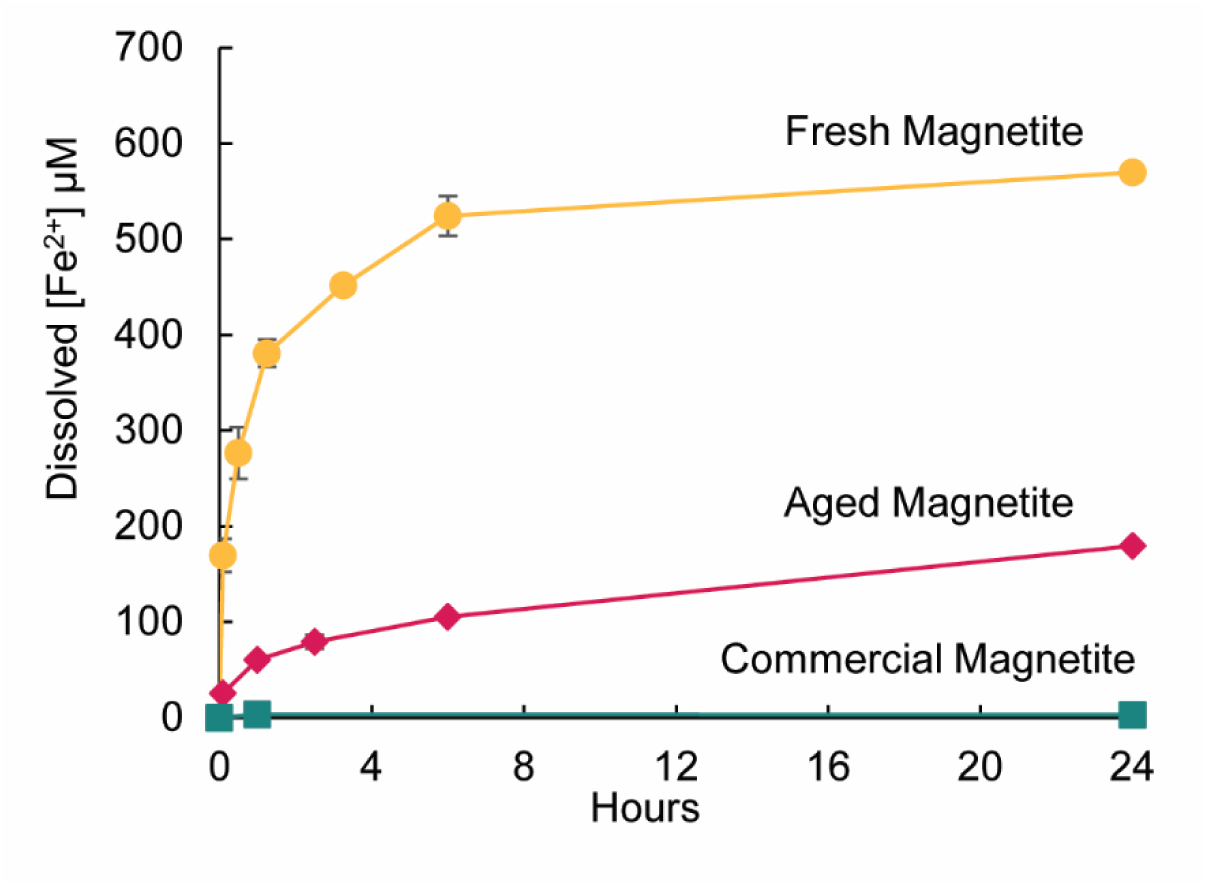
Dissolved Fe^2+^ released under anoxic conditions in 20 mM MES pH 6.0 from different magnetite types: fresh synthetic magnetite (gold; circles), aged synthetic magnetite (pink; diamonds), commercial magnetite (teal; squares). Error bars are ± one standard deviation of replicates.

### Sideroxydans lithotrophicus ES-1 growth on magnetite

Culturing experiments demonstrated that all magnetites supported growth of *S. lithotrophicus* ES-1. Over the course of a 14-day incubation, the cell numbers increased ∼50-fold in bottles containing all types of magnetite (Fig. 2; fresh magnetite 49.5×; aged magnetite 47.7×; commercial magnetite 59.4×). Cell numbers increased faster on the fresh and aged synthetic magnetites than on the commercial magnetite during the first four days, but at the end of the experiment, cell numbers were similar in all conditions (Fig. 2). The final cell yield of ∼2-3×10^7^ cells/mL is similar to the cell yield observed when *S. lithotrophicus* ES-1 was grown on 1 g/L of Fe(II)-smectite clay (7).

**Figure 2.**
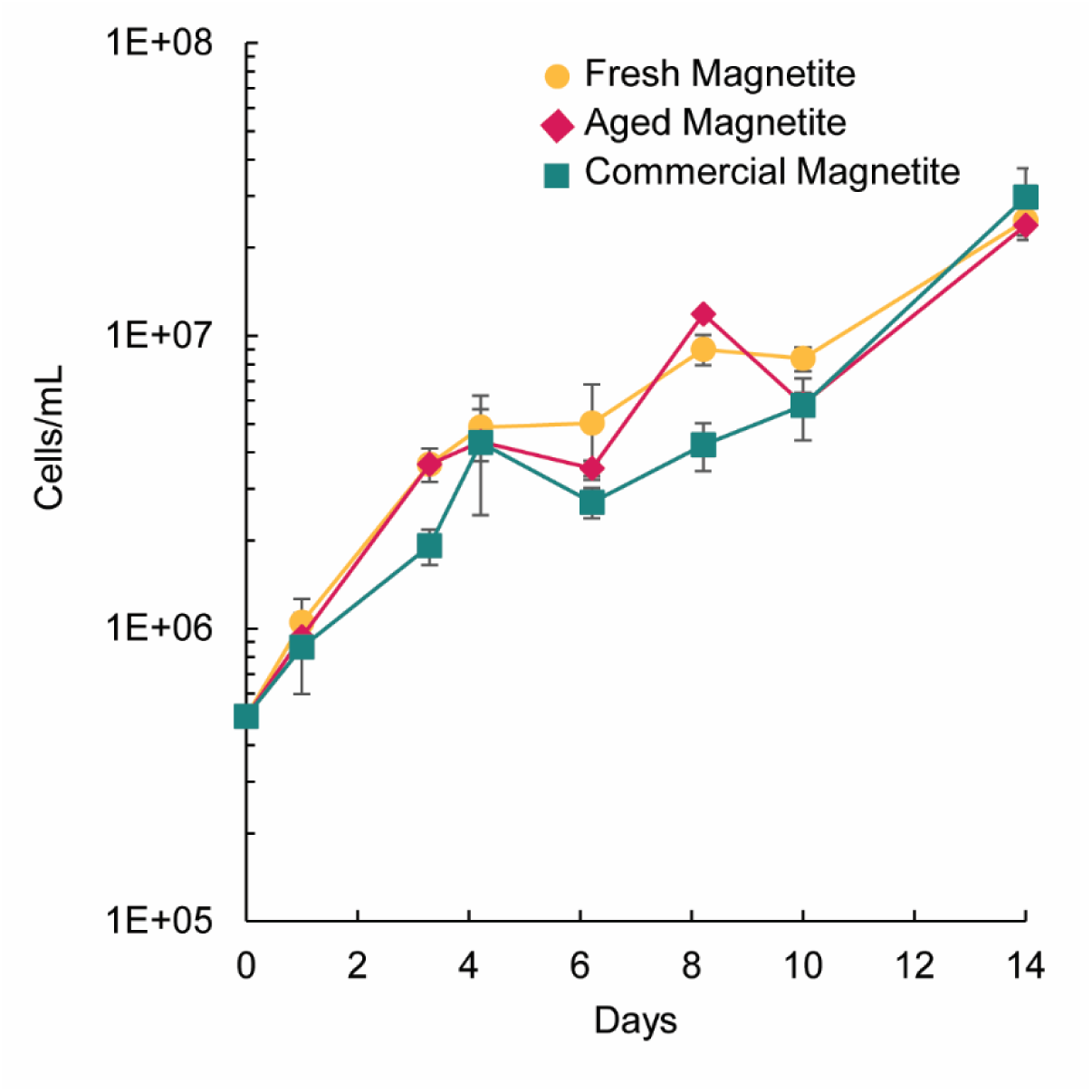
*S. lithotrophicus* ES-1 growth on fresh synthetic magnetite (gold; circles), aged synthetic magnetite (pink; diamonds), and commercial magnetite (teal; squares). Error bars are ± one standard deviation of replicates.

In previous experiments, we observed *S. lithotrophicus* ES-1 experienced exponential growth for five days with a maximum cell yield of <2×10^6^ cells/mL when provided with only 100 µM dissolved Fe^2+^ per day (in the form of Fe(II)-citrate) (17). In the magnetite cultures, *S. lithotrophicus* ES-1 reached more than one order of magnitude higher cell density and continued to build biomass through day 14 (Fig. 2), long after the available dissolved Fe^2+^ dropped below 100 µM (Fig. S2), suggesting *S. lithotrophicus* ES-1 is either promoting magnetite dissolution or accessing the solid magnetite directly.

### Dissolved and solid iron oxidation

We tracked the dissolved Fe^2+^ and Fe(II)/Fe(III) in magnetite to evaluate whether *S. lithotrophicus* ES-1 was oxidizing one or both forms of iron. In *S. lithotrophicus* ES-1 cultures with either fresh synthetic magnetite or aged synthetic magnetite, all dissolved Fe^2+^ was oxidized in the culture within three days (Fig. 3). The rate of oxidation of dissolved Fe^2+^ by oxygen was slower in the bottles without cells: in the fresh synthetic magnetite bottle, measurable dissolved Fe^2+^ remained at the end of the experiment while in the aged synthetic magnetite bottle, dissolved Fe^2+^ was measurable until day six. Because biotic oxidation is faster than the abiotic controls, *S. lithotrophicus* ES-1 cells are accelerating and/or catalyzing iron oxidation. In commercial magnetite bottles both with and without *S. lithotrophicus* ES-1, concentrations of dissolved Fe^2+^ were <10 μM at all time-points, suggesting that growth could be based primarily on oxidation of solid magnetite.

**Figure 3.**
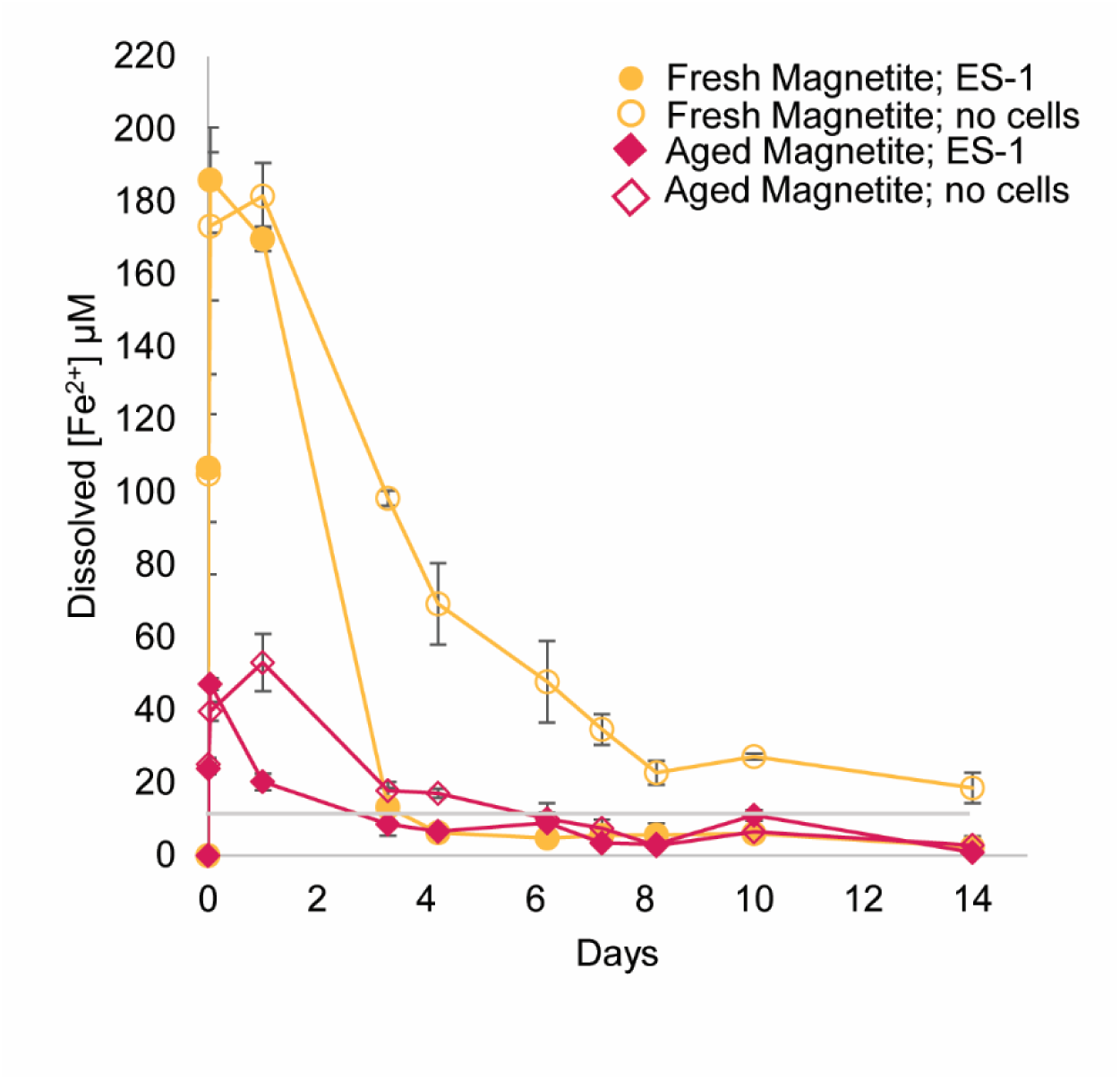
Dissolved Fe^2+^ remaining in cultures with *S. lithotrophicus* ES-1 (solid) or no cell controls (dashed; hollow) with different magnetite types: fresh synthetic magnetite (gold; circles), aged synthetic magnetite (pink; diamonds). Commercial magnetite measurements were always < 10 µM and were not plotted. Gray line at 10 µM is detection limit. Error bars are ± one standard deviation of replicates.

The Fe(II)/Fe(III) content of the magnetite was measured in minerals sampled over the course of the experiment (Fig. 4). At the start of the experiment, both of the synthetic magnetites were more reduced (Fe(II)/Fe(III) = 0.6-0.7) than stoichiometric magnetite (Fe(II)/Fe(III) = 0.5). In cultures with *S. lithotrophicus* ES-1, fresh synthetic magnetite was more oxidized (a lower Fe(II)/Fe(III) ratio) on day seven compared to the abiotic bottles (p<0.005); however, by the end of the experiment, ratios measured for both bottles were similar (Fig. 4A). In the aged synthetic magnetite bottles, there was more oxidation in the bottles with *S. lithotrophicus* ES-1 compared to bottles without cells starting at day 10 (p<0.005 at day 14; Fig. 4B). In contrast, for the commercial magnetite bottles, there was no difference in the ratio between bottles with cells and without cells. Despite the little measurable difference between rates of abiotic and biotic oxidation of solid magnetite, *S. lithotrophicus* ES-1 is capable of growing in the presence of solid magnetite as the sole electron donor. This suggests even if the cells do not accelerate the rate of oxidation as measured at the bulk level, they are still able to outcompete O_2_ for electrons. By day 14, the different types of magnetite were oxidized to a similar extent (Fe(II)/Fe(III) ∼ 0.4; Fig. 4), despite their various initial sizes, crystallinities, and starting Fe(II)/Fe(III) ratios. This suggests there is a proportion of Fe(II) in each of the magnetite structures that is inaccessible to either microbes or oxygen under these culture conditions.

**Figure 4.**
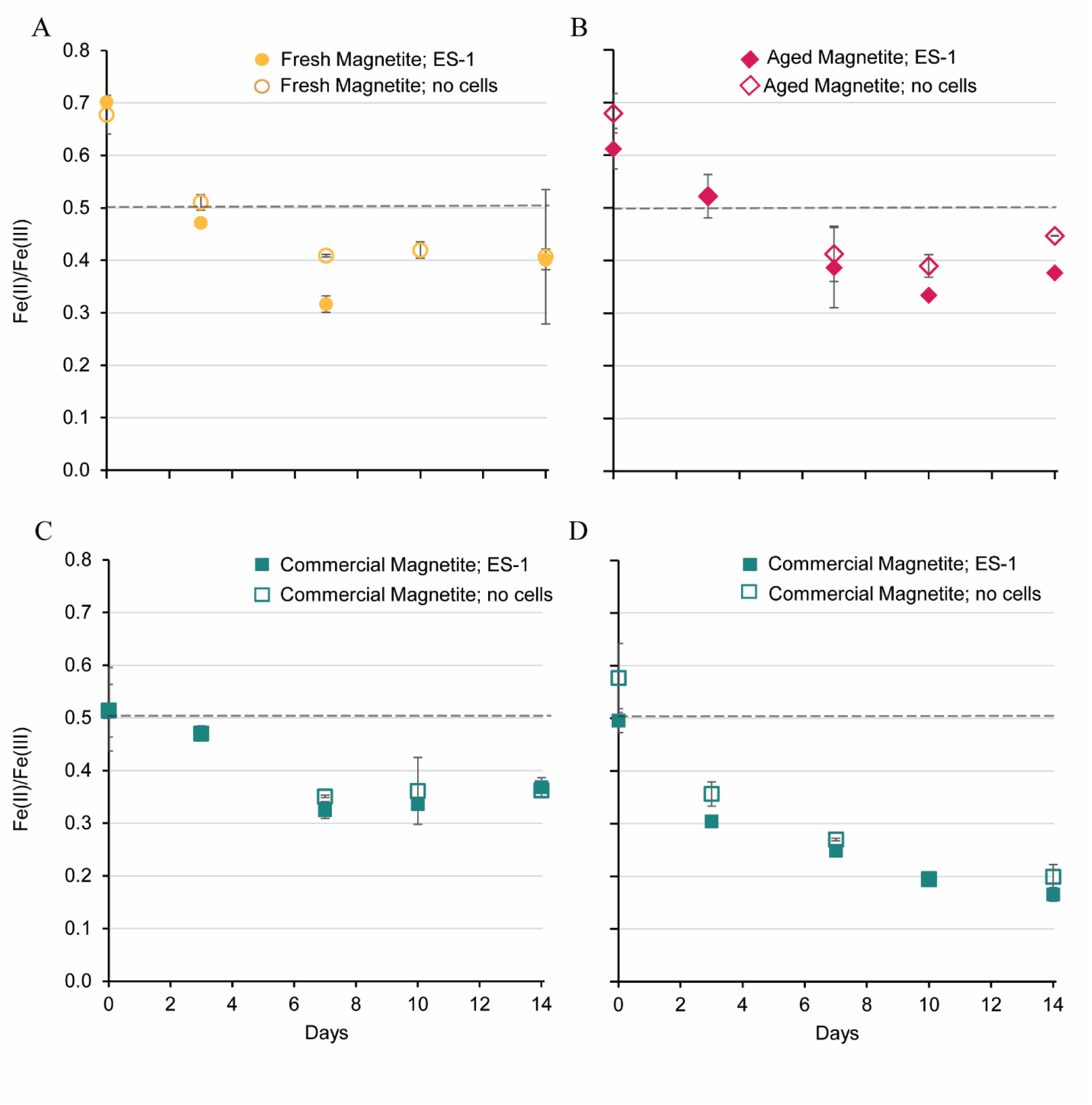
Measurements of the Fe(II) to Fe(III) ratio in acid-dissolved solid magnetite particles from cultures with *S. lithotrophicus* ES-1 (solid) or no cell controls (hollow) with different magnetite types: A) fresh synthetic magnetite; B) aged synthetic magnetite, C) commercial magnetite, full dissolution (6 M HCl; 24 hours), and D) commercial magnetite, partial dissolution (1 M HCl; 1 hour). Dashed line indicates the stoichiometric magnetite ratio. Error bars are ± one standard deviation of replicates.

Because these were bulk measurements, it was possible there was preferential oxidation of the surface that was obscured. To address this, the solid commercial magnetite particles were subjected to a partial dissolution step (∼15% dissolved in 1 M HCl) to measure reactive Fe(II) and Fe(III) of the surface. These results indicated that there was more oxidation of the surface (Fig. 4D) compared to the bulk particles (Fig. 4C), although there was still not much difference between cultures with *S. lithotrophicus* ES-1 and no-cell control bottles. Combining the results from all measurements of dissolved Fe^2+^ and Fe(II)/Fe(III) in magnetite suggests the dissolved Fe^2+^ is quickly oxidized by the microbes (Fig. 3) and the microbes could be concurrently accessing electrons from solid Fe(II) since the magnetite Fe(II)/Fe(III) ratio decreases by the first measurement on day 3 (Fig. 4).

### Proteome analyses

Quantitative proteomic analyses were performed to explore *S. lithotrophicus* ES-1 iron oxidation mechanisms on aqueous Fe(II) and magnetite. Magnetite cultures (Fig. 2) were compared to Fe(II)-citrate cultures (Fig. S3) at an early growth time-point (day 3 for the magnetites or day 2 for Fe(II)-citrate) or a late growth time-point (day 14 for magnetites or day 7 for Fe(II)-citrate). *S. lithotrophicus* grew faster on Fe(II)-citrate, but the cell number was normalized during collection for the proteome analysis. A total of 2309 out of 2978 proteins encoded in the genome (∼78%) were identified across all eight conditions (832-2068 proteins per sample), from 10-100 mL of culture, demonstrating the OFIC processing method used here is a significant improvement over the previous proteomic pipelines (7) for low biomass samples.

Principal component analyses (PCA) showed all of the Fe(II)-citrate grown samples were most similar to one another (Fig. 5A); these two time-points shared 93% of the proteins detected. There was a clear separation of the Fe(II)-citrate and magnetite samples along the component two axis (Fig. 5A), while the early and late time-point samples of the magnetites were separated along the component one axis. The magnetite samples showed more differentiation in the PCA, though all six magnetite samples did share 91% of the proteins detected, suggesting the magnetite-grown cultures express a core set of proteins. Together, these results show the growth phase and type of available Fe(II) source exert large influences on the variation within the protein expression profiles.

**Figure 5.**
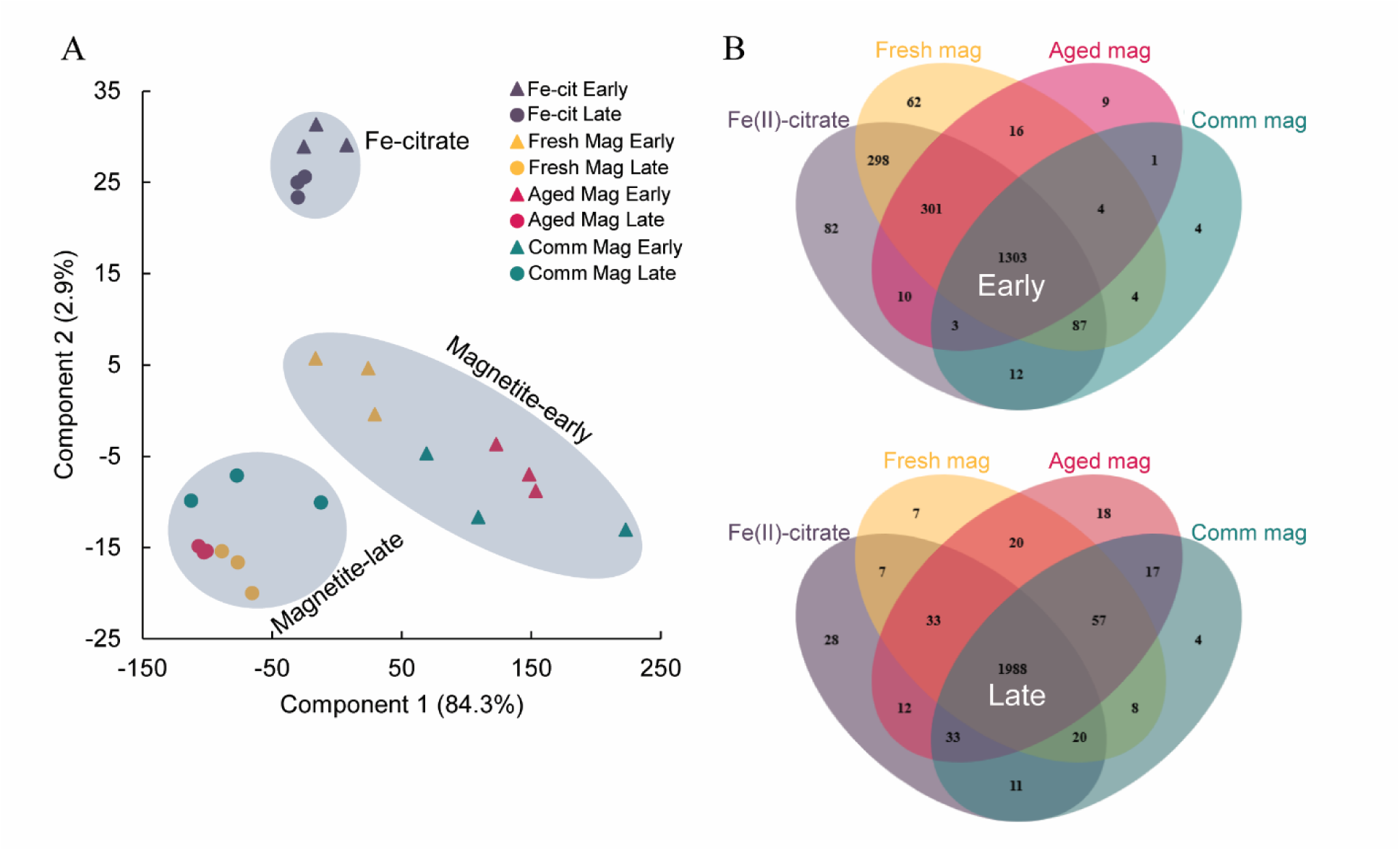
(A) Principal component analysis plot of the different sample types. Early time-points (triangles), late time-points (circles), Fe(II)-citrate (purple), fresh synthetic magnetite (gold), aged synthetic magnetite (pink), and commercial magnetite (teal). (B) Venn diagrams showing the number of proteins identified in each condition for the early time-point (top) and the late time-point (bottom). Fe-cit - Fe(II)-citrate; Mag – magnetite; Comm – commercial.

The fresh and aged synthetic magnetite cultures were more similar to the Fe(II)-citrate cultures at the early time-point. There were 298 proteins shared between the Fe(II)-citrate and fresh synthetic magnetite cultures that were not present in any other culture (Fig. 5B), and a pair-wise comparison found no statistically significant differences in protein abundances of the shared proteins between these two conditions. An additional 301 proteins were shared between the early time-point fresh synthetic, aged synthetic, and Fe(II)-citrate cultures. These cultures were the only ones with measurable amounts of dissolved Fe^2+^ (Fig. 3); thus, the shared proteins may represent mechanisms and adaptations for utilizing dissolved Fe^2+^.

*S. lithotrophicus* ES-1 encodes two distinct Fe(II) oxidation pathways, the MtoAB complex and Cyc2 (17, 34–36, 44). The Mto complex is comprised of a periplasmic decaheme MtoA (Slit_2497), and an outer membrane porin MtoB (Slit_2496) (35, 44). The same gene cluster encodes an inner membrane tetraheme protein CymA/ImoA (Slit_2495) and a periplasmic monoheme protein (MtoD; Slit_2498) (35, 53, 54). MtoA and CymA/ImoA were only detected in the magnetite cultures but not in the Fe(II)-citrate culture (Table 1) and were among the most differentially expressed proteins (Fig. 6; Table S1). The Mto complex was expressed even in the early magnetite cultures, suggesting the expression of Mto is controlled by the presence of the solid magnetite, regardless of the presence of dissolved Fe^2+^. These results are in agreement with the previous study with *S. lithotrophicus* ES-1 growing either on dissolved Fe(II)-citrate or solid Fe(II)-smectite clays, in which some of the proteins of the Mto complex were only detected in the solid Fe(II) cultures (7).

**Figure 6.**
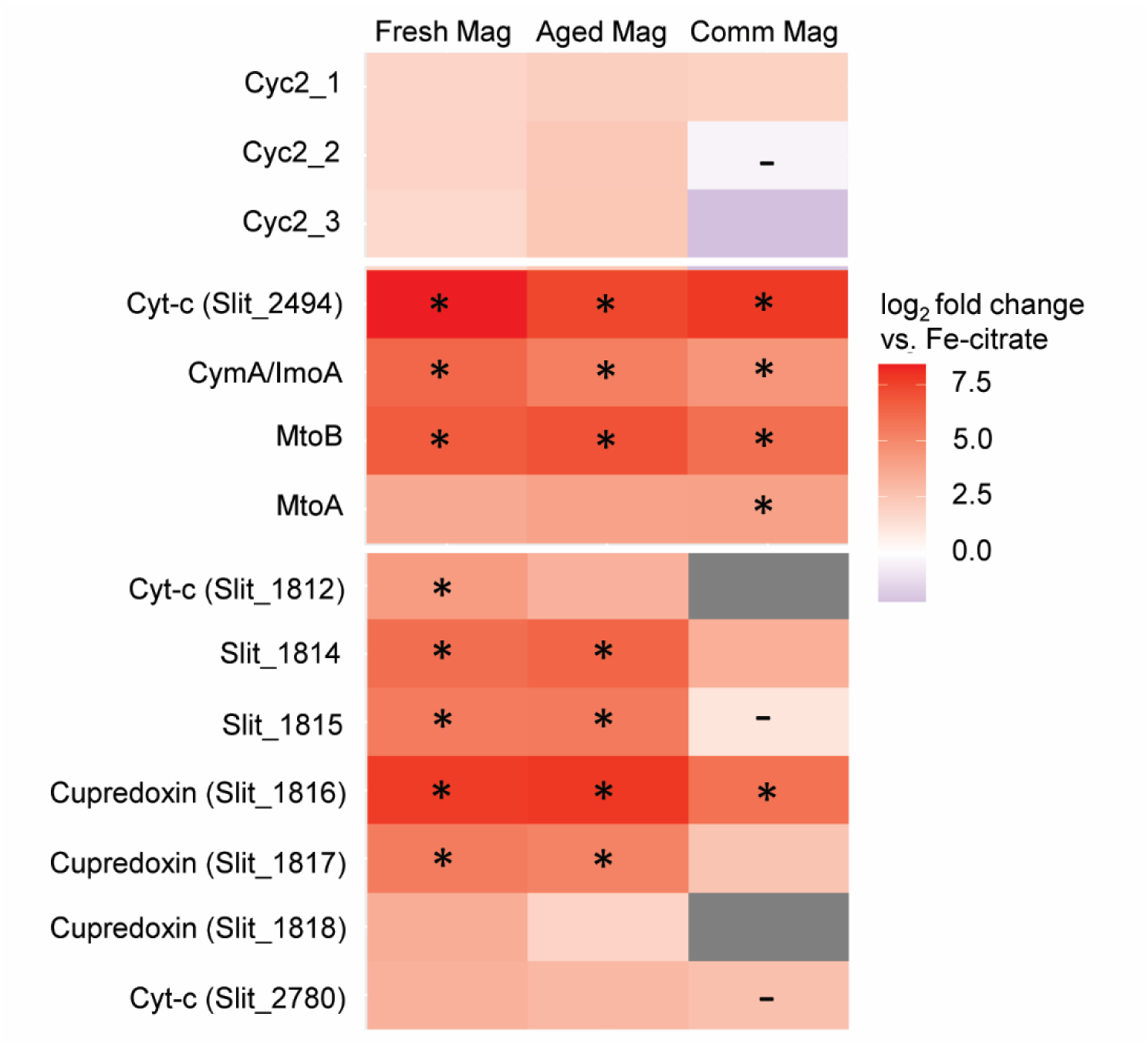
Heatmap of the log_2_ fold change in expression between each of the late magnetite samples and the late Fe(II)-citrate sample. Boxes marked with a (*) are within the top 1% of most differentially expressed proteins. All comparisons are statistically significant (P_adj_<0.05), with the exception of boxes marked with a (−). Gray boxes indicate protein was not detected in at least one comparison condition. Mag – magnetite; Comm - commercial

**Table 1.**
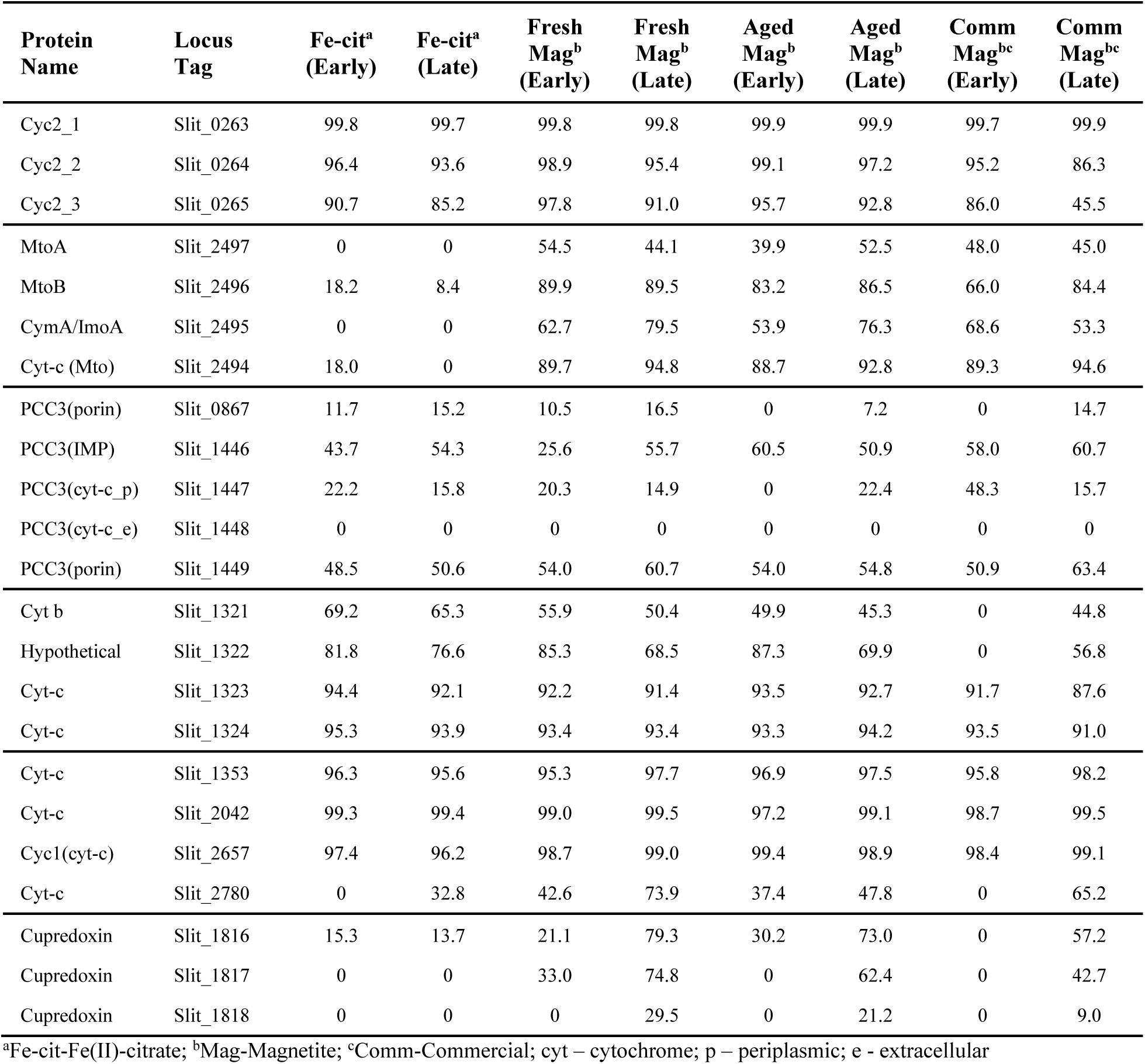
Maximum percentile of protein expression based on iBAQ values.

Expression of MtoD was not detected in any culture, but a protein encoded by the gene downstream of *cymA/imoA* (UniProt entry: D5CMP7, locus tag: Slit_2494) showed similar expression patterns as the other Mto-related proteins (Fig. 6; Table S1). This protein is poorly annotated but contains one heme binding motif and a transmembrane signal peptide, suggesting it could also be a periplasmic cytochrome involved in the Mto-based iron oxidation pathway.

The other Fe(II) oxidase in *S. lithotrophicus* ES-1 is the fused monoheme cytochrome porin, Cyc2 (17, 36, 37). *S. lithotrophicus* ES-1 has three isoforms of Cyc2, and all three were detected in both the Fe(II)-citrate and magnetite cultures. One isoform (Slit_0263; Cyc2_1) was one of the most highly expressed proteins in all samples (99.7th percentile; Table 1). The second isoform of the iron oxidase Cyc2 (Slit_0264) was one of the top expressed proteins in the early time-point samples of the fresh and aged synthetic magnetites. Cyc2 expression is high in all the conditions, even ones without measurable amounts of dissolved Fe(II). Together, the results show that while Cyc2 is highly expressed regardless of the presence of dissolved Fe^2+^, the Mto complex is only detected in the presence of solid Fe(II) substrates.

Other proteins that have been previously hypothesized to have a role in iron oxidation were also expressed. *S. lithotrophicus* ES-1 has two gene clusters encoding predicted porin-cytochrome complexes with multiheme cytochromes, with 18-28 CX(_2-4_)CH motifs, making them much larger than MtoA (34). The proteins encoded by the porin-cytochrome gene cluster Slit_0867-0870 were largely not detected. The proteins encoded by the second porin-cytochrome gene cluster Slit_1446-1449 were detected (with the exception of the predicted extracellular cytochrome, Slit_1448), and had similar levels of expression in all the samples (Table 1). Expression of the proteins encoded by the iron-responsive gene cluster (Slit_1321-1324) identified in Zhou *et al*. (17) were detected in all samples comparably, where Slit_1323 and Slit_1324 (predicted to be a monoheme and diheme cytochrome *c*, respectively) were expressed >87^th^ percentile (Table 1). The highly expressed and upregulated iron-responsive periplasmic cytochromes identified in Zhou *et al*. (17) (Slit_1353, Slit_2042, Slit_2657) were also highly abundant in all conditions here (>95th percentile; Table 1).

To identify additional proteins that could be specifically involved in solid Fe(II) oxidation, we compared the time-point with dominantly solid Fe(II) oxidation (late commercial magnetite) to one with solely dissolved Fe^2+^ oxidation (late Fe(II)-citrate). A cluster (Slit_1812-1818) of three cupredoxin-domain-containing proteins as well as a SCO1/SenC protein (Slit_1813), a predicted cytochrome (Slit_1812), and a few small hypothetical proteins (Slit_1814-1815) was more highly expressed in the late magnetite cultures compared to the Fe(II)-citrate cultures (Fig. 6; Table S1). Two of the cupredoxin-domain proteins have canonical multicopper oxidase motifs (55) (Slit_1817 and Slit_1818), the third does not (Slit_1816). The protein encoded by Slit_1816 was the most differentially expressed, with significantly higher expression in the late magnetite cultures compared to Fe(II)-citrate. We also identified a periplasmic cytochrome (Slit_2780) with higher expression in the late magnetite cultures than Fe(II)-citrate (Fig. 6; Table S1). These results suggest a possible role for these proteins in the oxidation of solid iron sources.

In all comparisons, there were more proteins with higher expression on the magnetites than on the Fe(II)-citrate samples. However, there were a number of proteins more highly expressed in the Fe(II)-citrate samples. Alternative complex III (Slit_0640-0646) is frequently implicated in iron oxidation pathways and was more highly expressed in the late Fe(II)-citrate cultures compared to the late commercial magnetite culture (Table S2). A cluster of proteins (Slit_0302-0307) were also more highly expressed in the Fe(II)-citrate compared to the late magnetite cultures (Table S2); these proteins may have a function in oxidative stress tolerance. Carbonic anhydrase (Slit_2956) and biotin synthase (BioB) were also more highly expressed in the Fe(II)-citrate cultures (Table S2), suggesting possible differences in CO_2_ metabolism. However, neither form I nor form II RuBisCo were more highly expressed on Fe(II)-citrate, and similar to the results found previously (17), form II was more highly expressed than form I in all cultures (Table S2).

## Discussion

Most of Earth’s iron is mineral-bound, potentially providing a vast source of energy if microbes can obtain electrons from minerals. Chemolithotrophic iron-oxidizing bacteria could hypothetically grow by mineral oxidation, but to date, there has been scant proof of this ability. Here we show that a well-studied iron oxidizer *Sideroxydans lithotrophicus* ES-1 can grow by oxidizing magnetite and constrain the likely enzymatic pathway via proteomics.

Unraveling iron oxidation mechanisms has been hampered by problems with culturing as well as RNA and protein extractions, but recent advances in on-filter, in-cell digestion proteomics methods now enable the study of proteins in low-yield, difficult to grow organisms like *S. lithotrophicus* ES-1 (46–48). Previous studies on smectite-grown *S. lithotrophicus* ES-1 included RT-qPCR and proteomics, but proteomics experiments were plagued with a number of issues including interference from iron and filter extractables, as well as the requirement for large volumes of culture to obtain enough cells (7). Proteomics (not transcriptomics) was required to unravel the mechanisms of magnetite oxidation by providing information on the presence of functional protein. Compared to previous studies, this study improved the overall protein detection rate (>78%) with fewer cells (∼3-4×10^8^ cells), and improved detection of multiheme cytochrome proteins, which enables us to better evaluate the mechanisms of iron oxidation. We are optimistic that the proteomics pipeline used here will enable the study of other organisms that are difficult to grow.

While it is well-known that iron-oxidizing bacteria like *S. lithotrophicus* ES-1 can grow by oxidizing dissolved Fe(II), here we established that it could grow on the solid mixed-valence iron mineral magnetite (Fe^II^Fe^III^_2_O_4_). We carefully followed the progression of iron redox in both dissolved and mineral phases and noted that magnetite oxidation occurred even in the presence of dissolved Fe(II), but the magnetite was never fully oxidized (lowest Fe(II):Fe(III) = 0.3; Fig. 4). Incomplete oxidation of magnetite was similarly observed in experiments with *Rhodopseudomonas palustris* TIE-1 and enrichment culture KS (1, 3), suggesting some amount of Fe(II) is not available for either biological or chemical oxidation. Partial oxidation may occur due to kinetic limitations in electron or iron atom diffusion through the magnetite structure (56), resulting in preferential oxidation of the surface as seen for the commercial magnetite (Fig. 4D). Another reason may be that the redox potential of magnetite increases with oxidation (56), causing some of the Fe(II) to be inaccessible due to thermodynamics. Given the various challenges with accessing minerals, it was a surprise that *S. lithotrophicus* ES-1 uses magnetite even in the presence of some dissolved Fe(II). Our results demonstrate that iron-oxidizing bacteria not only oxidize magnetite and dissolved Fe(II) individually, they can use both solutes and solids simultaneously, opening the question of how common such flexibility may be among other iron oxidizers.

The ability to access electrons from both dissolved and solid Fe(II) may require separate mechanisms appropriate for each type of Fe(II). Oxidation of solid electron donors likely involves the multiheme cytochrome-porin complex MtoAB, which could transfer electrons across the outer membrane. The MtoAB pathway seems to be specifically expressed by *S. lithotrophicus* ES-1 for solid Fe(II) oxidation. In previous studies, when *S. lithotrophicus* ES-1 was grown on Fe(II)-citrate, *mto* transcripts were very low (7, 17). In that study, which also analyzed incomplete proteomes, as well as in our current more comprehensive proteome work, most of the proteins of the Mto pathway were not detected during growth on dissolved Fe(II)-citrate (Table 1 and (7)). However, when grown on solid magnetite, MtoA/MtoB/CymA(ImoA)/Slit_2494 was one of the most significantly enriched sets of proteins (Fig. 6; Table S1), suggesting these proteins are reserved for oxidation of solid Fe(II) sources. This fits with previous research demonstrating that purified MtoA directly interacted with magnetite and was able to extract reactive Fe(II) from within solid ferrite spinel nanoparticles (57). The homologous MtrAB system of iron-reducing *Shewanella* has been shown to be electrochemically reversible and capable of electron uptake (58). Furthermore, the iron-oxidizing homolog in *Rhodopseudomonas palustris* TIE-1, PioAB, has been shown to play a role in the oxidation of solid electrodes (9, 43), and *R. palustris* can oxidize magnetite (although the proteins involved were not investigated) (1). Combined, these findings strongly imply that the MtoAB multiheme cytochrome-porin complex enables *S. lithotrophicus* to conduct extracellular electron uptake from solid electron donors (magnetite, smectite), and could do so in other organisms as well.

Dissolved Fe(II) is likely oxidized by another iron oxidase of *S. lithotrophicus* ES-1, Cyc2, which is also expressed during growth and oxidation of magnetite. Cyc2 is a monoheme cytochrome fused to a porin, and structural constraints lead to the prediction that Cyc2 must be an oxidase of aqueous Fe^2+^ ions (36). Previous work showed Cyc2, specifically the first isoform of Cyc2 (Slit_0263), is highly expressed in all growth conditions, including dissolved Fe(II)-citrate, solid Fe(II)-smectite clays, and thiosulfate (7, 17). That continues to be true during growth on magnetite. We hypothesize that dissolved Fe(II) (i.e. Fe^2+^) is the preferred electron donor of *S. lithotrophicus* ES-1 and thus it maintains readiness to oxidize dissolved Fe(II) regardless of its presence by constitutively expressing Cyc2 at high levels. It is also possible that dissolved Fe(II) at concentrations below our detection limit is being shed from the magnetite, and is acting as an electron shuttle, prompting the expression of Cyc2. Another possibility could be that Cyc2 plays a role in iron sensing. In any case, *S. lithotrophicus* ES-1 appears to express its iron oxidases differently. The smaller monoheme cytochrome Cyc2 is expressed under all conditions, while the larger and more energetically expensive complex MtoAB is only expressed when necessary, i.e. when a solid electron source is present.

Magnetite oxidation may also involve copper-containing proteins, cupredoxins. Three uncharacterized copper-containing proteins were significantly more highly expressed in the magnetite cultures than the Fe(II)-citrate cultures (Fig. 6; Table S1). Two of these proteins possess typical multicopper oxidase motifs (Slit_1817 and Slit_1818). The third (Slit_1816) does not and is significantly larger, with additional domains similar to adhesions and polysaccharide lyases. These proteins are encoded together in the genome, along with a few smaller proteins, a SCO1/SenC protein, and a hypothetical cytochrome. Similar gene clusters are found in other organisms, mostly other members of the Burkholderiales like *Paraburkholderia* and *Ralstonia* but also in *Anaeromyxobacter* and *Steroidobacteraceae* (Fig. S4). While the roles of these cupredoxin proteins are not known, they are predicted to be extracellular proteins, which would enable access to magnetite particles. Other copper-containing proteins have been reported with ferroxidase (59) and Mn(II)-oxidase activity (60, 61), and play a role in iron oxidation in acidophilic iron oxidizers (62, 63); thus it is plausible that the cupredoxins identified here are playing a role in solid Fe(II) oxidation in *S. lithotrophicus* ES-1.

Given the growing recognition of microbial mineral oxidation, it will be important to increase our understanding of the mechanisms in order to recognize and trace the activities of mineral-oxidizing microbes. The evidence obtained so far suggests that magnetite and Fe(II)-smectite oxidation in *S. lithotrophicus* ES-1 involves the MtoAB complex, a decaheme cytochrome-porin complex homologous to the *Shewanella* Fe-reductase MtrAB. In studies of *Shewanella, Geobacter,* and other FeRB, we have learned that multiheme cytochromes (MHCs) are well-suited for redox interactions with minerals (64, 65), and their useful characteristics translate well into advantages for oxidizing minerals: 1) When housed in an outer membrane porin, MHCs can transfer electrons across the membrane to or from a mineral. An outer membrane-embedded MHC could either have direct contact with a mineral or transfer to/from extracellular MHC that contact minerals. 2) Unlike single heme cytochromes, MHCs have wide ranges of redox potentials that overlap with mineral redox potentials, which also span wide ranges and can change as minerals are oxidized and reduced. The multiheme cytochrome MtoA exhibits a range of redox potentials (−400 mV to +100 mV vs. SHE; (44, 66) that overlaps with 10-20 nm magnetite (−480 to +50 mV vs. SHE) and smectites (e.g., −600 to +0 mV for SWa-1; −400 to +400 mV for SWy-2; (56, 67, 68)). 3) MHCs can act as capacitors to store electrons, enabling microbes to continue making energy if there is an interruption in electron supply. This is more likely for minerals, which may be periodically exhausted of electron supply, in contrast to dissolved substrates that tend to be in more constant supply. So, overall, although MHCs are resource intensive – MtoAB is larger than Cyc2 (1165 vs ∼440 amino acids, ten heme cofactors vs one) - the investment in biosynthetic energy and resources would enable access to electrons stored in redox-active sedimentary minerals.

The utility of MHCs to FeOB is suggested by the number and diversity of MHCs in known FeOB. More than 60% of the iron-oxidizing Gallionellaceae possess a putative decaheme or larger cytochrome, suggesting many of these iron oxidizers may be able to utilize solid electron donors (45). Many of the Gallionellaceae possess multiple MHC gene clusters. For instance, *S. lithotrophicus* ES-1 encodes MtoAB plus at least two other MHC complexes known as PCC3; these other cytochromes could be used by *S. lithotrophicus* ES-1 to oxidize different solid substrates. It remains to be seen if these are deployed under different conditions individually for distinct substrates or work simultaneously. It will be necessary to further constrain the functional relationships between specific MHCs, minerals, and growth conditions to enable gene- and protein-based tracking of microbial mineral oxidation. Multiheme cytochromes are being increasingly recognized in diverse organisms (69–74), opening the possibility of discovering new mineral-oxidizing organisms and broadening our understanding of the functionality of multiheme cytochromes.

Overall, our work expands our understanding of how magnetite can promote microbial growth, which has implications for biogeochemical cycling in sediments, aquifers, and rock-hosted environments. In these systems, magnetite can serve as an electron donor to microbes, but then can be re-reduced by iron/mineral-reducing microbes. Once recharged, the magnetite can be discharged again by FeOB, and so on, cycling back and forth, making magnetite a biogeochemical redox buffer that also supports growth and associated C, N, and P transformations. As we increasingly recognize the metabolic flexibility and adaptability of iron oxidizers like *S. lithotrophicus* ES-1 to the varied iron sources on Earth, this will help us understand the active role of iron oxidizers in iron mineral biogeochemical cycling throughout the Earth’s environments.

## Experimental Methods

### Magnetite synthesis and preparations

Synthetic magnetite (Fe_3_O_4_) was prepared according to the protocol outlined in Byrne *et al*. (1). Solutions of 1 M FeCl_2_ and 2 M FeCl_3_ were prepared in anoxic 0.3 M HCl. The two solutions were combined and added dropwise into anoxic NaOH (25%) with continuous stirring at 870 rpm in an anaerobic chamber. The black precipitate was collected and washed with anoxic water to remove residual chloride ions. The synthetic magnetite was dried in a desiccator chamber within the anaerobic chamber, then ground with a mortar and pestle under anoxic conditions. This preparation was the fresh synthetic magnetite. For the aged synthetic magnetite, fresh synthetic magnetite was resuspended in anoxic water adjusted to ∼pH 10.0 with NaOH. This solution was heated to 95 °C for one week in a sealed serum bottle in a water bath, then autoclaved for 30 min. at 121 °C. The aged synthetic magnetite was returned to the anaerobic chamber, then dried and ground as described above. Commercial magnetite (Iron(II,III) oxide; Cas. No. 1317-61-9) was purchased from Sigma-Aldrich (Cat. No. 637106). All types of magnetite were sterilized in an autoclave for 30 min. at 121 °C as a dry powder under anoxic conditions before use.

### Magnetite characterization

The mineral phase of all dry, sterilized products was confirmed by X-ray diffraction (XRD) using a Bruker D8 Powder XRD with Cu Kα radiation. To avoid oxidation during data collection, the samples were loaded into quartz capillary tubes (Charlessupper; outside dimension 1.0 mm) and sealed with silicone in the anaerobic chamber. Data was obtained from 10-70° 2θ with a step size of 0.05 and an acquisition time of 1s. The data was collected on autorepeat for at least 15 hours to enhance the diffraction signal. The raw spectra were processed with background subtraction and matched against the database in the DIFFRAC.EVA program. Particle size was calculated based on the Scherrer equation using the full width at half maximum (FWHM) of the six highest intensity peaks (75).

Dissolved and solid Fe measurements were collected using a modified 1,10-phenanthroline assay (76, 77). At selected time points, samples were taken, then centrifuged in the anaerobic chamber at 13000 × *g* for 5 min. The supernatant was collected to measure the dissolved Fe^2+^. The precipitates were fully dissolved in 6 M HCl or partially dissolved in 1 M HCl under anoxic conditions for 24 h to determine the ferrous to ferric ratio (Fe(II)/Fe(III)) of the bulk mineral or of the mineral surface (78, 79), respectively. The solutions were diluted 1:4 (1 M HCl) or 1:40 (6 M HCl) with anoxic water. For Fe(II) measurements, 20 µL samples were mixed with 80 µL anoxic water, 50 µL 0.1% 1,10-phenanthroline, and 50 µL 3 M sodium acetate, pH 5.5 in the anaerobic chamber. After a 15-minute incubation, absorbance was measured at 512 nm and compared to a standard curve. For total Fe measurements, 80 µL 10% hydroxylamine hydrochloride was used instead of water, and the sample was incubated for 1 hour before addition of the phenanthroline reagent and acetate buffer. Significant differences were determined using a two-tailed, paired *t*-test with a cutoff threshold of 0.05.

### Cultures

*Sideroxydans lithotrophicus* ES-1 was pre-grown in modified Wolfe’s minimal medium (MWMM) plus trace minerals and vitamins (17, 80, 81), buffered with 20 mM 2-(N-morpholino)ethanesulfonic acid (MES) pH 6.0 with 10 mM thiosulfate as the electron donor and 2% oxygen as the electron acceptor. Thiosulfate was chosen for the pre-cultures to avoid introducing Fe(III) into experimental reactors. During experiments, synthetic or commercial magnetite (1 g/L; 12.9 mM Fe) with 2% oxygen was utilized as the electron donor and electron acceptor, respectively. The headspace was flushed daily with 2% oxygen/20% carbon dioxide/78% nitrogen. *S. lithotrophicus* ES-1 was also grown in MWMM with 20 mM MES pH 6.0, 5 mM citrate, 2% oxygen headspace, and daily additions of 200 μM FeCl_2_. The cell number was determined by counting SYTO 13-stained cells under fluorescent microscopy using a Petroff-Hausser counting chamber.

### Proteomics

For each sample type, ∼ 3-4×10^8^ cells (10-100 mL) were processed following the on-filter in-cell digestion protocol described previously (46). In brief, culture was loaded 3 mL at a time onto the cartridges, then centrifuged at 500 x *g* for 1 min. After collecting the total number of cells on the cartridges, cells were incubated with pure methanol at 4 °C for 30 min. Afterwards, the cartridges were spun to discard the methanol, and the proteins were reduced and alkylated, digested with trypsin (Promega, Madison, WI), eluted, then desalted using C18-based StageTips (CDS Analytical, Oxford, PA) as described previously (46). The LC-MS/MS analysis was performed using an Ultimate 3000 RSLCnano system and Orbitrap Eclipse mass spectrometer installed with FAIMS Pro Interface (ThermoScientific) also as described previously (46).

For proteome quantitation, raw MS data were processed using MaxQuant (82) and Andromeda software suite (version 2.4.2.0). The protein database of *Sideroxydans lithotrophicus* ES-1 (taxonomy_id:580332; 2,978 protein sequences) was downloaded from the UniProtKB website (https://www.uniprot.org/). The enzyme specificity was set to ‘Trypsin’; variable modifications include oxidation of methionine, and acetyl (protein N-terminus); fixed modification includes carbamidomethylation of cysteine. The maximum missed cleavage sites were set to 2 and the minimum number of amino acids required for peptide identification was 7. The false discovery rate (FDR) was set to 1% for protein and peptide identifications. MaxLFQ function embedded in MaxQuant was enabled for label-free quantitation, and the LFQ minimum ratio count was set to 1. Proteins identified as reverse hits, potential contaminants, or only by site-modification were filtered out from the “proteinGroups.txt” output file. The LFQ values were log_2_ transformed, filtered by at least two valid values out of three replicates in at least one group, and imputed using the default “normal distribution” method in Perseus (version 2.0.6.0) (83).

### Analysis tools

Venn diagrams were created using goodcalculators.com/venn-diagram-maker. Heat maps were made in R using ggplot2 (84). Gene cluster comparisons were performed using cblaster and clinker https://cagecat.bioinformatics.nl/ (85, 86). Protein subcellular localization predicted using (PSORTb v3.0.3 https://www.psort.org/psortb/ (87). All the statistical analyses, including the Student’s T-test with Permutation-based FDR, were performed using Perseus (version 2.0.6.0) (83).

## Data availability

The MS raw files associated with this study have been deposited to the MassIVE server (https://massive.ucsd.edu/) with the dataset identifier MSV000093770, and is publicly available as of the date of submission. Unprocessed protein intensities and iBAQ values (Table S3) and processed pairwise comparisons (Table S4) are available as part of the supplement.

## Acknowledgements

This research was funded by the National Science Foundation (BIO-1817651). Access to the University of Delaware Mass Spectrometry Core was supported by the Institutional Development Award (IDeA) from the National Institute of Health’s National Institute of General Medical Sciences under grant number P20GM103446. The University of Delaware Mass Spectrometry Facility is supported by The National Institutes of Health Center of Biomedical Research Excellence (NIH-COBRE) Program, with a grant from the National Institute of General Medical Sciences (P20GM104316). We thank the Delaware Biotechnology Institute (DBI) for core instrumentation, Gerald Poirier in the Advanced Materials Characterization Laboratory at the University of Delaware for help with XRD, Katherine Martin and Papa Nii Asare-Okai in the University of Delaware Mass Spectrometry Facility, and James Byrne for helpful discussions. Competing interests: The author(s) declare none. The content is solely the responsibility of the authors and does not necessarily represent the official views of the National Institutes of Health.

## Notes

### Competing Interest Statement

The authors have declared no competing interest.

### Summary of Updates

Minor edits to the introduction and discussion, clarification of dissolved and solid iron oxidation, updated Fig. 6

## References

1. Byrne JM, Klueglein N, Pearce C, Rosso KM, Appel E, Kappler A. 2015. Redox cycling of Fe(II) and Fe(III) in magnetite by Fe-metabolizing bacteria. Science 347:1473–1476.

2. Chakraborty A, Roden EE, Schieber J, Picardal F. 2011. Enhanced Growth of Acidovorax sp. Strain 2AN during Nitrate-Dependent Fe(II) Oxidation in Batch and Continuous-Flow Systems▿. 24. Appl Environ Microbiol 77:8548–8556.

3. Weber KA, Picardal FW, Roden EE. 2001. Microbially Catalyzed Nitrate-Dependent Oxidation of Biogenic Solid-Phase Fe(II) Compounds. Environ Sci Technol 35:1644–1650.

4. Miot J, Li J, Benzerara K, Sougrati MT, Ona-Nguema G, Bernard S, Jumas J-C, Guyot F. 2014. Formation of single domain magnetite by green rust oxidation promoted by microbial anaerobic nitrate-dependent iron oxidation. Geochimica et Cosmochimica Acta 139:327–343.

5. Percak-Dennett E, He S, Converse B, Konishi H, Xu H, Corcoran A, Noguera D, Chan C, Bhattacharyya A, Borch T, Boyd E, Roden EE. 2017. Microbial acceleration of aerobic pyrite oxidation at circumneutral pH. Geobiology 15:690–703.

6. Benzine J, Shelobolina E, Xiong MY, Kennedy DW, McKinley JP, Lin X, Roden E. 2013. Fe-phyllosilicate redox cycling organisms from a redox transition zone in Hanford 300 Area sediments. Front Microbiol 4.

7. Zhou N, Kupper RJ, Catalano JG, Thompson A, Chan CS. 2022. Biological Oxidation of Fe(II)-Bearing Smectite by Microaerophilic Iron Oxidizer Sideroxydans lithotrophicus Using Dual Mto and Cyc2 Iron Oxidation Pathways. Environ Sci Technol 56:17443–17453.

8. Shelobolina ES, Konishi H, Xu H, Benzine J, Xiong MY, Wu T, Blöthe M, Roden E. 2012. Isolation of Phyllosilicate–Iron Redox Cycling Microorganisms from an Illite–Smectite Rich Hydromorphic Soil. Front Microbiol 3.

9. Bose A, Gardel EJ, Vidoudez C, Parra EA, Girguis PR. 2014. Electron uptake by iron-oxidizing phototrophic bacteria. Nat Commun 5:3391.

10. Rabaey K, Rodríguez J, Blackall LL, Keller J, Gross P, Batstone D, Verstraete W, Nealson KH. 2007. Microbial ecology meets electrochemistry: electricity-driven and driving communities. The ISME Journal 1:9–18.

11. Eddie BJ, Wang Z, Malanoski AP, Hall RJ, Oh SD, Heiner C, Lin B, Strycharz-Glaven SMY 2016. ‘Candidatus Tenderia electrophaga’, an uncultivated electroautotroph from a biocathode enrichment. International Journal of Systematic and Evolutionary Microbiology 66:2178–2185.

12. Rowe AR, Chellamuthu P, Lam B, Okamoto A, Nealson KH. 2015. Marine sediments microbes capable of electrode oxidation as a surrogate for lithotrophic insoluble substrate metabolism. Frontiers in Microbiology 5.

13. Gupta D, Guzman MS, Bose A. 2020. Extracellular electron uptake by autotrophic microbes: physiological, ecological, and evolutionary implications. Journal of Industrial Microbiology and Biotechnology 47:863–876.

14. Comolli LR, Luef B, Chan CS. 2011. High-resolution 2D and 3D cryo-TEM reveals structural adaptations of two stalk-forming bacteria to an Fe-oxidizing lifestyle. Environ Microbiol 13:2915–2929.

15. Chan CS, Fakra SC, Emerson D, Fleming EJ, Edwards KJ. 2011. Lithotrophic iron-oxidizing bacteria produce organic stalks to control mineral growth: implications for biosignature formation. The ISME Journal 5:717–727.

16. Emerson D, Moyer C. 1997. Isolation and characterization of novel iron-oxidizing bacteria that grow at circumneutral pH. 12. Appl Environ Microbiol 63:4784–4792.

17. Zhou N, Keffer JL, Polson SW, Chan CS. 2022. Unraveling Fe(II)-Oxidizing Mechanisms in a Facultative Fe(II) Oxidizer, Sideroxydans lithotrophicus Strain ES-1, via Culturing, Transcriptomics, and Reverse Transcription-Quantitative PCR. Applied and Environmental Microbiology 88:e01595–21.

18. Zhou N, Luther GW, Chan CS. 2021. Ligand effects in abiotic and biotic Fe(II) oxidation by the microaerophile *Sideroxydans lithotrophicus*. Environmental Science & Technology 10.1021/acs.est.1c00497.

19. Fortney NW, He S, Converse BJ, Boyd ES, Roden EE. 2018. Investigating the Composition and Metabolic Potential of Microbial Communities in Chocolate Pots Hot Springs. Frontiers in Microbiology 9.

20. Eze MO, Lütgert SA, Neubauer H, Balouri A, Kraft AA, Sieven A, Daniel R, Wemheuer B. 2020. Metagenome Assembly and Metagenome-Assembled Genome Sequences from a Historical Oil Field Located in Wietze, Germany. Microbiol Resour Announc 9:e00333–20.

21. Bethencourt L, Bochet O, Farasin J, Aquilina L, Borgne TL, Quaiser A, Biget M, Michon-Coudouel S, Labasque T, Dufresne A. 2020. Genome reconstruction reveals distinct assemblages of *Gallionellaceae* in surface and subsurface redox transition zones. FEMS Microbiology Ecology 96:fiaa036.

22. Zhang S-Y, Su J-Q, Sun G-X, Yang Y, Zhao Y, Ding J, Chen Y-S, Shen Y, Zhu G, Rensing C, Zhu Y-G. 2017. Land scale biogeography of arsenic biotransformation genes in estuarine wetland: Microbial biogeography of As biotransformation genes. 6. Environ Microbiol 19:2468–2482.

23. Parks DH, Rinke C, Chuvochina M, Chaumeil P-A, Woodcroft BJ, Evans PN, Hugenholtz P, Tyson GW. 2017. Recovery of nearly 8,000 metagenome-assembled genomes substantially expands the tree of life. 11. Nature Microbiology 2:1533–1542.

24. Buck M, Garcia SL, Fernandez L, Martin G, Martinez-Rodriguez GA, Saarenheimo J, Zopfi J, Bertilsson S, Peura S. 2021. Comprehensive dataset of shotgun metagenomes from oxygen stratified freshwater lakes and ponds. 1. Sci Data 8:131.

25. Tian R, Ning D, He Z, Zhang P, Spencer SJ, Gao S, Shi W, Wu L, Zhang Y, Yang Y, Adams BG, Rocha AM, Detienne BL, Lowe KA, Joyner DC, Klingeman DM, Arkin AP, Fields MW, Hazen TC, Stahl DA, Alm EJ, Zhou J. 2020. Small and mighty: adaptation of superphylum Patescibacteria to groundwater environment drives their genome simplicity. Microbiome 8:51.

26. Poghosyan L, Koch H, Frank J, van Kessel MAHJ, Cremers G, van Alen T, Jetten MSM, Op den Camp HJM, Lücker S. 2020. Metagenomic profiling of ammonia- and methane-oxidizing microorganisms in two sequential rapid sand filters. Water Res 185:116288.

27. Bell E, Lamminmäki T, Alneberg J, Andersson AF, Qian C, Xiong W, Hettich RL, Frutschi M, Bernier-Latmani R. 2020. Active sulfur cycling in the terrestrial deep subsurface. ISME J 14:1260–1272.

28. Wrighton KC, Thomas BC, Sharon I, Miller CS, Castelle CJ, VerBerkmoes NC, Wilkins MJ, Hettich RL, Lipton MS, Williams KH, Long PE, Banfield JF. 2012. Fermentation, Hydrogen, and Sulfur Metabolism in Multiple Uncultivated Bacterial Phyla. Science 337:1661–1665.

29. Ahmed M, Saup CM, Wilkins MJ, Lin L-S. 2020. Continuous ferric iron-dosed anaerobic wastewater treatment: Treatment performance, sludge characteristics, and microbial composition. Journal of Environmental Chemical Engineering 8:103537.

30. Ceballos-Escalera A, Pous N, Chiluiza-Ramos P, Korth B, Harnisch F, Bañeras L, Balaguer MD, Puig S. 2021. Electro-bioremediation of nitrate and arsenite polluted groundwater. Water Research 190:116748.

31. Chan CS, Dykes GE, Hoover RL, Limmer MA, Seyfferth AL. 2023. Gallionellaceae in rice root plaque: metabolic roles in iron oxidation, nutrient cycling, and plant interactions. Applied and Environmental Microbiology 89:e00570–23.

32. Cooper RE, Wegner C-E, McAllister SM, Shevchenko O, Chan CS, Küsel K. 2020. Draft genome sequence of *Sideroxydans* sp. strain CL21, an Fe(II)-oxidizing bacterium. Microbiol Resour Announc 9:e01444–19.

33. Woodcroft BJ, Singleton CM, Boyd JA, Evans PN, Emerson JB, Zayed AAF, Hoelzle RD, Lamberton TO, McCalley CK, Hodgkins SB, Wilson RM, Purvine SO, Nicora CD, Li C, Frolking S, Chanton JP, Crill PM, Saleska SR, Rich VI, Tyson GW. 2018. Genome-centric view of carbon processing in thawing permafrost. 7716. Nature 560:49–54.

34. He S, Barco RA, Emerson D, Roden EE. 2017. Comparative genomic analysis of neutrophilic iron(II) oxidizer genomes for candidate genes in extracellular electron transfer. Front Microbiol 8:1584.

35. Emerson D, Field EK, Chertkov O, Davenport KW, Goodwin L, Munk C, Nolan M, Woyke T. 2013. Comparative genomics of freshwater Fe-oxidizing bacteria: implications for physiology, ecology, and systematics. Front Microbiol 4:254.

36. Keffer JL, McAllister SM, Garber AI, Hallahan BJ, Sutherland MC, Rozovsky S, Chan CS. 2021. Iron Oxidation by a Fused Cytochrome-Porin Common to Diverse Iron-Oxidizing Bacteria. mBio 12:e01074–21.

37. McAllister SM, Polson SW, Butterfield DA, Glazer BT, Sylvan JB, Chan CS. 2020. Validating the Cyc2 neutrophilic iron oxidation pathway using meta-omics of Zetaproteobacteria iron mats at marine hydrothermal vents. mSystems 5:e00553–19.

38. Jiao Y, Newman DK. 2007. The pio operon is essential for phototrophic Fe(II) oxidation in Rhodopseudomonas palustris TIE-1. JB 189:1765–1773.

39. Beliaev AS, Saffarini DA. 1998. Shewanella putrefaciens mtrB Encodes an Outer Membrane Protein Required for Fe(III) and Mn(IV) Reduction. J Bacteriol 180:6292–6297.

40. Pitts KE, Dobbin PS, Reyes-Ramirez F, Thomson AJ, Richardson DJ, Seward HE. 2003. Characterization of the Shewanella oneidensis MR-1 Decaheme Cytochrome MtrA: EXPRESSION IN ESCHERICHIA COLI CONFERS THE ABILITY TO REDUCE SOLUBLE FE(III) CHELATES *. Journal of Biological Chemistry 278:27758–27765.

41. Richardson DJ, Butt JN, Fredrickson JK, Zachara JM, Shi L, Edwards MJ, White G, Baiden N, Gates AJ, Marritt SJ, Clarke TA. 2012. The ‘porin–cytochrome’ model for microbe-to-mineral electron transfer. Molecular Microbiology 85:201–212.

42. White GF, Edwards MJ, Gomez-Perez L, Richardson DJ, Butt JN, Clarke TA. 2016. Chapter three - mechanisms of bacterial extracellular electron exchange, p. 87–138. In Poole, RK (ed.), Advances in Microbial Physiology. Academic Press.

43. Gupta D, Sutherland MC, Rengasamy K, Meacham JM, Kranz RG, Bose A. 2019. Photoferrotrophs produce a PioAB electron conduit for extracellular electron uptake. mBio 10.

44. Liu J, Wang Z, Belchik SM, Edwards MJ, Liu C, Kennedy DW, Merkley ED, Lipton MS, Butt JN, Richardson DJ, Zachara JM, Fredrickson JK, Rosso KM, Shi L. 2012. Identification and characterization of MtoA: A decaheme c-type cytochrome of the neutrophilic Fe(II)-oxidizing bacterium *Sideroxydans lithotrophicus* ES-1. Front Microbiol 3:37.

45. Hoover RL, Keffer JL, Polson SW, Chan CS. 2023. Gallionellaceae pangenomic analysis reveals insight into phylogeny, metabolic flexibility, and iron oxidation mechanisms. mSystems 8:e00038–23.

46. Martin KR, Le HT, Abdelgawad A, Yang C, Lu G, Keffer JL, Zhang X, Zhuang Z, Asare-Okai PN, Chan CS, Batish M, Yu Y. 2024. Development of an efficient, effective, and economical technology for proteome analysis. Cell Reports Methods 4:100796.

47. Kelly V, Al-Rawi A, Lewis D, Kustatscher G, Ly T. 2022. Low Cell Number Proteomic Analysis Using In-Cell Protease Digests Reveals a Robust Signature for Cell Cycle State Classification. Mol Cell Proteomics 21:100169.

48. Hatano A, Takami T, Matsumoto M. 2023. In situ digestion of alcohol-fixed cells for quantitative proteomics. J Biochem 173:243–254.

49. Cornell RM, Schwertmann U. 2003. The Iron Oxides: Structure, Properties, Reactions, Occurences and Uses, 2nd ed. Wiley-VCH Verlag GmbH & Co. KGaA.

50. Byrne JM, van der Laan G, Figueroa AI, Qafoku O, Wang C, Pearce CI, Jackson M, Feinberg J, Rosso KM, Kappler A. 2016. Size dependent microbial oxidation and reduction of magnetite nano- and micro-particles. 1. Sci Rep 6:30969.

51. Cartledge BT, Marcotte AR, Herckes P, Anbar AD, Majestic BJ. 2015. The Impact of Particle Size, Relative Humidity, and Sulfur Dioxide on Iron Solubility in Simulated Atmospheric Marine Aerosols. Environ Sci Technol 49:7179–7187.

52. Thompson A, Chadwick OA, Rancourt DG, Chorover J. 2006. Iron-oxide crystallinity increases during soil redox oscillations. Geochimica et Cosmochimica Acta 70:1710–1727.

53. Beckwith CR, Edwards MJ, Lawes M, Shi L, Butt JN, Richardson DJ, Clarke TA. 2015. Characterization of MtoD from *Sideroxydans lithotrophicus*: a cytochrome c electron shuttle used in lithoautotrophic growth. Front Microbiol 6.

54. Jain A, Coelho A, Madjarov J, Paquete CM, Gralnick JA. 2022. Evidence for Quinol Oxidation Activity of ImoA, a Novel NapC/NirT Family Protein from the Neutrophilic Fe(II)-Oxidizing Bacterium Sideroxydans lithotrophicus ES-1. mBio 13:e02150–22.

55. Gräff M, Buchholz PCF, Le Roes-Hill M, Pleiss J. 2020. Multicopper oxidases: modular structure, sequence space, and evolutionary relationships. Proteins 88:1329–1339.

56. Gorski CA, Nurmi JT, Tratnyek PG, Hofstetter TB, Scherer MM. 2010. Redox Behavior of Magnetite: Implications for Contaminant Reduction. Environ Sci Technol 44:55–60.

57. Liu J, Pearce CI, Liu C, Wang Z, Shi L, Arenholz E, Rosso KM. 2013. Fe3–xTixO4 Nanoparticles as Tunable Probes of Microbial Metal Oxidation. J Am Chem Soc 135:8896–8907.

58. Ross DE, Flynn JM, Baron DB, Gralnick JA, Bond DR. 2011. Towards Electrosynthesis in Shewanella: Energetics of Reversing the Mtr Pathway for Reductive Metabolism. PLOS ONE 6:e16649.

59. Huston WM, Jennings MP, McEwan AG. 2002. The multicopper oxidase of Pseudomonas aeruginosa is a ferroxidase with a central role in iron acquisition. Molecular Microbiology 45:1741–1750.

60. Romano CA, Zhou M, Song Y, Wysocki VH, Dohnalkova AC, Kovarik L, Paša-Tolić L, Tebo BM. 2017. Biogenic manganese oxide nanoparticle formation by a multimeric multicopper oxidase Mnx. Nat Commun 8:746.

61. Soldatova AV, Butterfield C, Oyerinde OF, Tebo BM, Spiro TG. 2012. Multicopper Oxidase Involvement in Both Mn(II) and Mn(III) Oxidation during Bacterial Formation of MnO2. J Biol Inorg Chem 17:1151–1158.

62. Cox JC, Boxer DH. 1978. The purification and some properties of rusticyanin, a blue copper protein involved in iron(II) oxidation from Thiobacillus ferro-oxidans. Biochemical Journal 174:497–502.

63. Appia-Ayme C, Guiliani N, Ratouchniak J, Bonnefoy V. 1999. Characterization of an Operon Encoding Two c-Type Cytochromes, an aa3-Type Cytochrome Oxidase, and Rusticyanin in Thiobacillus ferrooxidans ATCC 33020. Appl Environ Microbiol 65:4781–4787.

64. Edwards MJ, Richardson DJ, Paquete CM, Clarke TA. 2020. Role of multiheme cytochromes involved in extracellular anaerobic respiration in bacteria. Protein Science 29:830–842.

65. Shi L, Squier TC, Zachara JM, Fredrickson JK. 2007. Respiration of metal (hydr)oxides by Shewanella and Geobacter: a key role for multihaem c-type cytochromes. Molecular Microbiology 65:12–20.

66. Paquete CM, Morgado L, Salgueiro CA, Louro RO. 2022. Molecular Mechanisms of Microbial Extracellular Electron Transfer: The Importance of Multiheme Cytochromes. 6. FBL 27:174.

67. Kappler A, Bryce C, Mansor M, Lueder U, Byrne JM, Swanner ED. 2021. An evolving view on biogeochemical cycling of iron. Nat Rev Microbiol 19:360–374.

68. Gorski CA, Klüpfel LE, Voegelin A, Sander M, Hofstetter TB. 2013. Redox Properties of Structural Fe in Clay Minerals: 3. Relationships between Smectite Redox and Structural Properties. Environ Sci Technol 47:13477–13485.

69. Salgueiro CA, Morgado L, Silva MA, Ferreira MR, Fernandes TM, Portela PC. 2022. From iron to bacterial electroconductive filaments: Exploring cytochrome diversity using *Geobacter* bacteria. Coordination Chemistry Reviews 452:214284.

70. Deng X, Dohmae N, Nealson KH, Hashimoto K, Okamoto A. 2018. Multi-heme cytochromes provide a pathway for survival in energy-limited environments. Science Advances 4:eaao5682.

71. Carlson HK, Iavarone AT, Gorur A, Yeo BS, Tran R, Melnyk RA, Mathies RA, Auer M, Coates JD. 2012. Surface multiheme c-type cytochromes from Thermincola potens and implications for respiratory metal reduction by Gram-positive bacteria. Proceedings of the National Academy of Sciences 109:1702–1707.

72. Thomas SH, Wagner RD, Arakaki AK, Skolnick J, Kirby JR, Shimkets LJ, Sanford RA, Löffler FE. 2008. The Mosaic Genome of Anaeromyxobacter dehalogenans Strain 2CP-C Suggests an Aerobic Common Ancestor to the Delta-Proteobacteria. PLoS One 3:e2103.

73. Downing BE, Gupta D, Nayak DD. 2023. The dual role of a multi-heme cytochrome in methanogenesis: MmcA is important for energy conservation and carbon metabolism in Methanosarcina acetivorans. Molecular Microbiology 119:350–363.

74. Zhang X, Joyce GH, Leu AO, Zhao J, Rabiee H, Virdis B, Tyson GW, Yuan Z, McIlroy SJ, Hu S. 2023. Multi-heme cytochrome-mediated extracellular electron transfer by the anaerobic methanotroph ‘Candidatus Methanoperedens nitroreducens.’ Nat Commun 14:6118.

75. Patterson AL. 1939. The Scherrer Formula for X-Ray Particle Size Determination. Phys Rev 56:978–982.

76. Tarafder PK, Thakur R. 2013. An Optimised 1,10-Phenanthroline Method for the Determination of Ferrous and Ferric Oxides in Silicate Rocks, Soils and Minerals. Geostandards and Geoanalytical Research 37:155–168.

77. de Mello Gabriel GV, Pitombo LM, Rosa LMT, Navarrete AA, Botero WG, do Carmo JB, de Oliveira LC. 2021. The environmental importance of iron speciation in soils: evaluation of classic methodologies. Environ Monit Assess 193:63.

78. Wallmann K, Hennies K, König I, Petersen W, Knauth H-D. 1993. New procedure for determining reactive Fe(III) and Fe(II) minerals in sediments. Limnology and Oceanography 38:1803–1812.

79. Porsch K, Kappler A. 2011. FeII oxidation by molecular O2 during HCl extraction. Environ Chem 8:190–197.

80. Emerson D, Merrill Floyd M. 2005. Enrichment and Isolation of Iron-Oxidizing Bacteria at Neutral pH, p. 112–123. In Methods in Enzymology. Academic Press.

81. Hädrich A, Taillefert M, Akob DM, Cooper RE, Litzba U, Wagner FE, Nietzsche S, Ciobota V, Rösch P, Popp J, Küsel K. 2019. Microbial Fe(II) oxidation by Sideroxydans lithotrophicus ES-1 in the presence of Schlöppnerbrunnen fen-derived humic acids. FEMS Microbiology Ecology 95:fiz034.

82. Tyanova S, Temu T, Cox J. 2016. The MaxQuant computational platform for mass spectrometry-based shotgun proteomics. Nat Protoc 11:2301–2319.

83. Tyanova S, Temu T, Sinitcyn P, Carlson A, Hein MY, Geiger T, Mann M, Cox J. 2016. The Perseus computational platform for comprehensive analysis of (prote)omics data. 9. Nat Methods 13:731–740.

84. Wickham H. 2009. ggplot2: Elegant Graphics for Data Analysis. Springer-Verlag, New York, NY.

85. Gilchrist CLM, Booth TJ, van Wersch B, van Grieken L, Medema MH, Chooi Y-H. 2021. cblaster: a remote search tool for rapid identification and visualization of homologous gene clusters. Bioinformatics Advances 1:vbab016.

86. Gilchrist CLM, Chooi Y-H. 2021. clinker & clustermap.js: automatic generation of gene cluster comparison figures. Bioinformatics 37:2473–2475.

87. Yu NY, Wagner JR, Laird MR, Melli G, Rey S, Lo R, Dao P, Sahinalp SC, Ester M, Foster LJ, Brinkman FSL. 2010. PSORTb 3.0: improved protein subcellular localization prediction with refined localization subcategories and predictive capabilities for all prokaryotes. Bioinformatics 26:1608–1615.

