## Supplement for "Microbial magnetite oxidation via MtoAB porin-multiheme cytochrome complex in *Sideroxydans lithotrophicus* ES-1"

List of Figures and Tables:

Figure S1 – page 2

Figure S2 – page 3

Figure S3 – page 4

Figure S4 – page 5

Table S1 – page 6

Table S2 – page 7

Table S3 – Unprocessed LQF intensities and iBAQ values (xlsx file)

Table S4 – Processed pairwise comparisons (xlsx file)

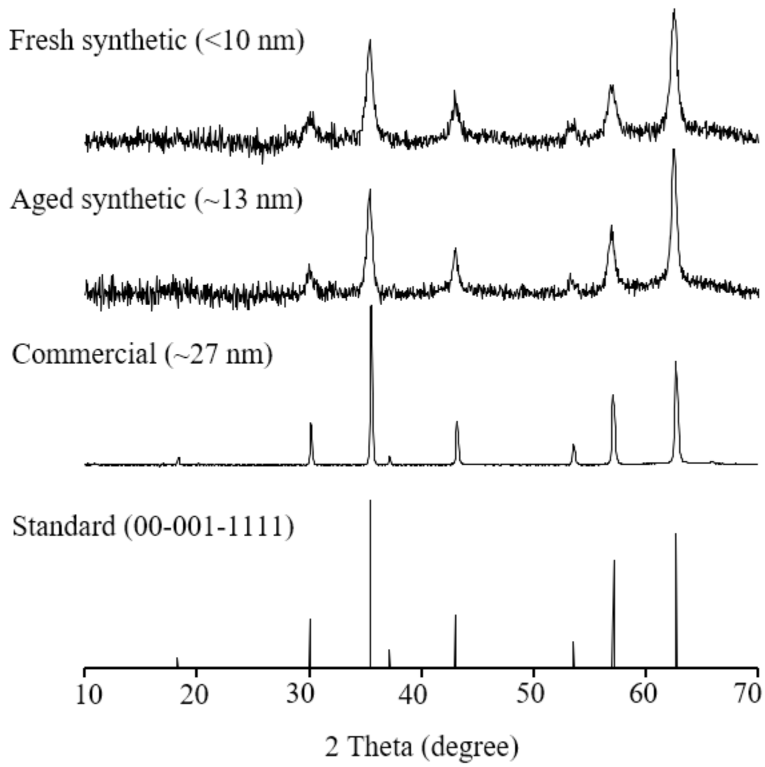


**Figure S1**. XRD patterns of (from top to bottom): fresh synthetic magnetite, aged synthetic magnetite, commercial magnetite, and magnetite ($Fe_3O_4$) standard 00-001-1111. Calculated particles sizes in parentheses.

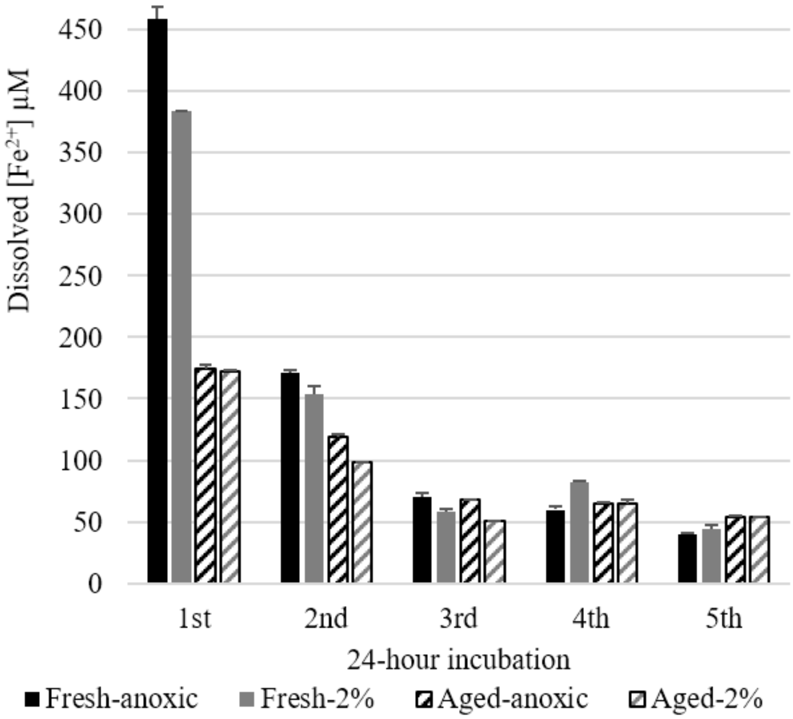


**Figure S2**. Dissolved $Fe^{2+}$ released under anoxic (black) or 2% oxygen (gray) conditions in 20 mM MES pH 6.0 from different magnetite types: fresh synthetic magnetite (solid bars), aged synthetic magnetite (hashed bars). Commercial magnetite was not measured since its first release is below the limit of detection (<10 µM). Error bars are + one standard deviation for replicates.

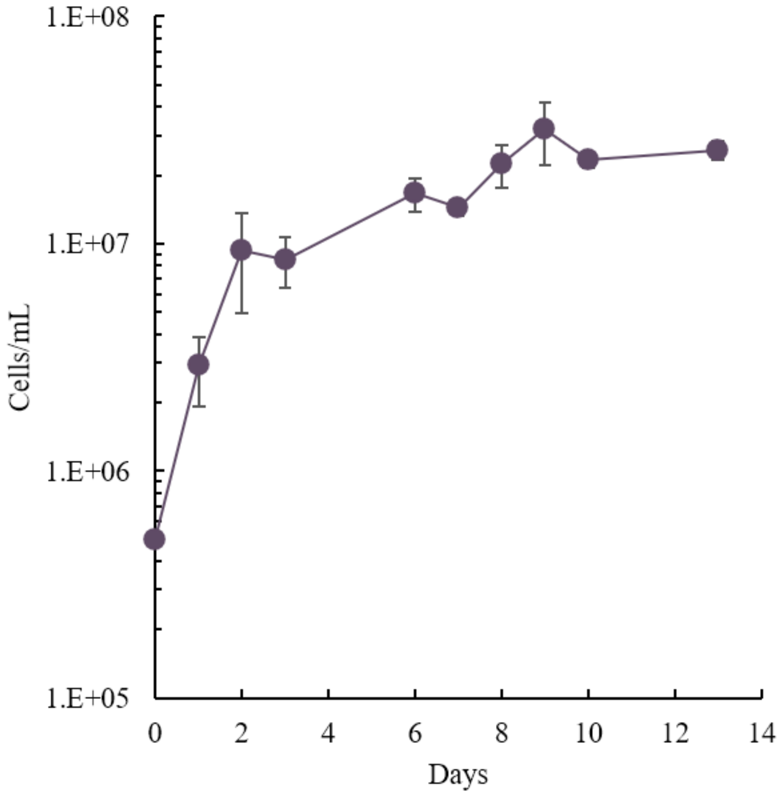


**Figure S3**. *S. lithotrophicus* ES-1 growth on Fe(II)-citrate (200 µM/day). Error bars are ± one standard deviation for replicates.

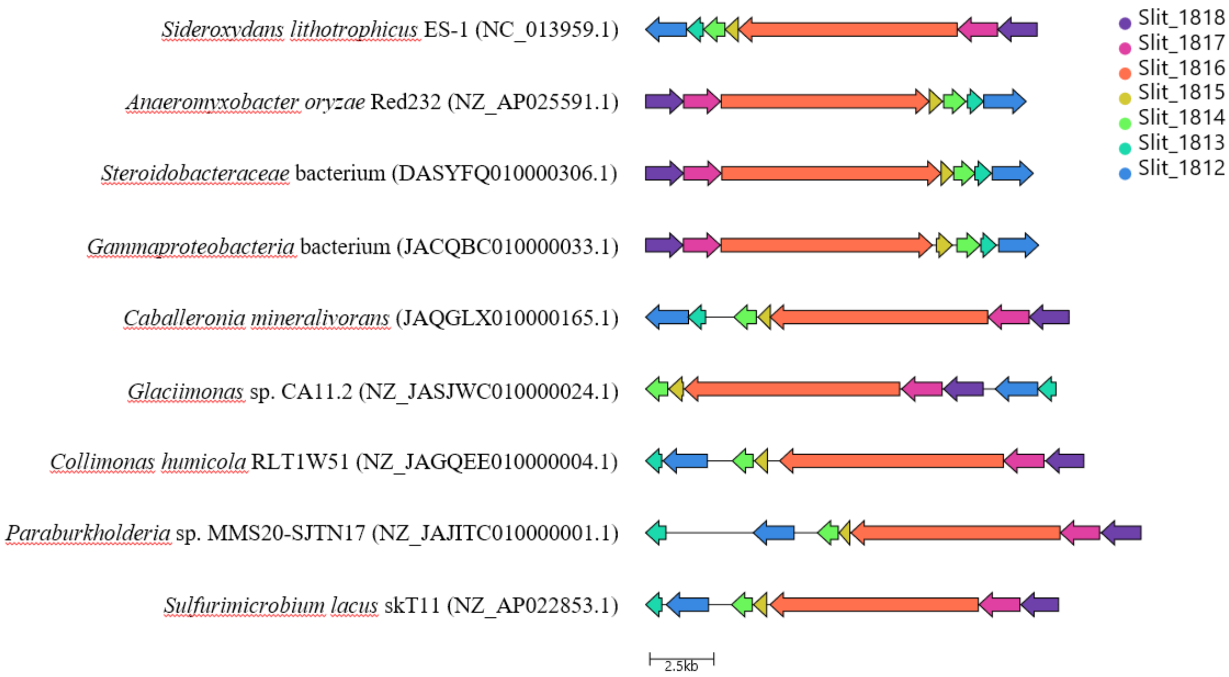


**Figure S4**. Gene cluster comparison between *Sideroxydans lithotrophicus* ES-1 and other selected organisms possessing a similar three cupredoxin-containing cluster. Accession number in parentheses. Figure generated using cblaster and clinker (https://cagecat.bioinformatics.nl).

**Table S1.** p-values for each $log_2$ fold change presented in Figure 6.

| Protein | Fresh Mag | Aged Mag | Comm Mag |
|---|---|---|---|
| Cyc2_1 | 0.0006 | 0.00006 | 0.03 |
| Cyc2_2 | 0.0007 | 0.00006 | 0.5 |
| Cyc2_3 | 0.008 | 0.0001 | 0.03 |
| Cyt-c (Slit_2494) | 0.000007 | 0.000002 | 0.0005 |
| CymA/ImoA | 0.00002 | 0.00005 | 0.02 |
| MtoB | 0.0007 | 0.001 | 0.005 |
| MtoA | 0.0005 | 0.001 | 0.0008 |
| Slit_1812 | 0.00004 | 0.0005 | |
| Slit_1814 | 0.00003 | 0.000003 | 0.02 |
| Slit_1815 | 0.0002 | 0.0000006 | 0.3 |
| Slit_1816 | 0.00008 | 0.00003 | 0.001 |
| Slit_1817 | 0.0003 | 0.0003 | 0.03 |
| Slit_1818 | 0.004 | 0.001 | |
| Slit_2780 | 0.001 | 0.0008 | 0.02 |

Mag – magnetite; Comm – commerical; Cyt - cytochrome

**Table S2.** Maximum percentile of protein expression based on iBAQ values and $log_2$ fold change for late Fe(II)-citrate cultures compared to commercial magnetite.

| Protein Name | Locus Tag | Fe-cit (Late) | Fresh Mag (Late) | Aged Mag (Late) | Comm Mag (Late) | Log2 Foldchange (FC/CM) | p-value |
|---|---|---|---|---|---|---|---|
| ACIII | Slit_0640 | 83.5 | 79.9 | 83.4 | 21 | 2.53876 | 0.004 |
| ACIII | Slit_0641 | 95.1 | 93.8 | 95.1 | 74.3 | 2.20014 | 0.02 |
| ACIII | Slit_0642 | 94.8 | 91.3 | 94.5 | 72.7 | 2.30073 | 0.03 |
| ACIII | Slit_0643 | 74.8 | 62 | 52.2 | 27.5 | 3.22756 | 0.004 |
| ACIII | Slit_0644 | 96.4 | 95.1 | 97.4 | 76.4 | 2.21245 | 0.02 |
| ACIII | Slit_0645 | 78.4 | 73.1 | 74.7 | 36.8 | 2.16156 | 0.04 |
| Rubrerythrin | Slit_0302 | 99.9 | 92.9 | 93.3 | 94.4 | 3.65931 | 0.005 |
| Uncharacterized | Slit_0303 | 86.4 | 38.9 | 38.7 | 41.2 | 2.70022 | 0.006 |
| Uncharacterized | Slit_0304 | 93.2 | 13.1 | 41.6 | 19.1 | 6.11879 | 0.003 |
| Ferritin family | Slit_0305 | 99.4 | 87.5 | 91.7 | 88.2 | 2.98962 | 0.01 |
| NnrS family | Slit_0307 | 45 | 0 | 6.7 | 0 | 1.31085 | 0.05 |
| Carbonic anhydrase | Slit_2956 | 84.7 | 0 | 0 | 0 | 4.99871 | 0.0007 |
| Biotin synthase | bioB | 71.6 | 13.6 | 13.2 | 13.3 | 3.31605 | 0.007 |
| cbbM | Slit_0022 | 97.6 | 96.7 | 98 | 97.5 | -0.56375 | 0.4 |
| cbbL | cbbL | 10.1 | 7.6 | 0 | 12.2 | -1.18568 | 0.4 |
| cbbS | Slit_0986 | 10.3 | 0 | 0 | 30.8 | -2.00883 | 0.04 |

Fe-cit – Fe(II)-citrate; Mag – magnetite; Comm – commercial; FC – late Fe(II)-citrate; CM – late commerical magnetite
